# Out-of-register parallel β-sheets and antiparallel β-sheets coexist in 150 kDa oligomers formed by Aβ(1-42)

**DOI:** 10.1101/2020.03.03.974394

**Authors:** Yuan Gao, Cong Guo, Jens O. Watzlawik, Elizabeth J. Lee, Danting Huang, Huan-Xiang Zhou, Terrone L. Rosenberry, Anant K. Paravastu

**Author notes:** Correspondence should be addressed to: Anant K. Paravastu, Associate Professor, School of Chemical and Biomolecular Engineering, Georgia Institute of Technology, 311 Ferst Drive NW, Atlanta, GA 30332-0100.

## Abstract

We present solid-state NMR measurements of β-strand secondary structure and inter-strand organization within a 150 kDa oligomeric aggregate of the 42-residue variant of the Alzheimer’s amyloid-β peptide (Aβ(1-42)). This oligomer is characterized by a structure that cannot be explained by any previously proposed model for aggregated Aβ. We build upon our previous report of a β-strand spanned by residues 30-42, which arranges into an antiparallel β-sheet. New results presented here indicate that there is a second β-strand formed by residues 11-24. We show negative results for NMR experiments designed to reveal antiparallel β-sheets formed by this β-strand. Remarkably, we show that this strand is organized into a parallel β-sheet despite the co-existence of an antiparallel β-sheet in the same structure. In addition, the in-register parallel β-sheet commonly observed for amyloid fibril structure does not apply to residues 11-24 in the 150 kDa oligomer. Rather, we present evidence for an inter-strand registry shift of 3 residues that alternates in direction between adjacent molecules along the β-sheet. We corroborated this unexpected scheme for β-strand organization using multiple 2-dimensional NMR and ^13^C-^13^C dipolar recoupling experiments. Our findings indicate a previously unknown assembly pathway and inspire a suggestion as to why this aggregate does not grow to larger sizes.

## Introduction

Research on the role of amyloid-β (Aβ) peptide assembly in Alzheimer’s disease (AD) can be described in terms of two presently unanswered questions: 1) what aggregated Aβ structures are possible under different aggregation pathways; and 2) what roles do different aggregated Aβ structures play in cellular pathology in the brain? The evidence for the central role played by Aβ aggregation in AD is reviewed elsewhere [1]. However, aggregation pathways are complex, producing multiple possible structures broadly classified as oligomers (dimers through 50-mers), protofibrils (hundreds of molecules), and amyloid fibrils (thousands to millions of molecules). The documented ability of the 40-residue variant of Aβ (Aβ(1-40)) to form multiple distinct fibril structures [2-4] highlights the complexity of Aβ aggregation pathways and suggests that there may also be considerable diversity in possible oligomer and protofibril structures. Recently reported structures of fibrils of the 42-residue variant of Aβ (Aβ(1-42)) [5-9] and mutant Aβ(1-40) [10, 11] differs from any previously reported structures of Aβ(1-40) fibrils, indicating that a small difference in primary structure could dramatically affect aggregation pathways. Furthermore, the toxicity of Aβ aggregates has been shown to be structure dependent: investigations of Aβ toxicity in cell culture, animal models, and human brain tissue has highlighted the special toxicity of oligomers over toxicity levels associated with protofibrils and fibrils [12]. In vivo, oligomers could diffuse over longer length scales than the larger Aβ aggregates and may interact in specific ways with cellular membranes and receptors. Furthermore, Aβ oligomers comprise an extremely diverse group [13] and include endogenous [14] as well as synthetic species prepared in vitro [15].

From a biophysical perspective, it is a mystery why Aβ would assemble into any low-molecular-weight oligomeric structure without undergoing further assembly into amyloid fibrils. Measurements based on methods such as Fourier transform infrared spectroscopy (FTIR) and circular dichroism (CD) establish β-strand secondary structure for oligomers. In general, peptides in β-strand conformations readily undergo assembly into β-sheets as backbone hydrogen bond donors and acceptors pair with complementary groups of neighboring peptide backbones. A β-sheet of any size would have dangling hydrogen bond donors and acceptors at its ends, and these ends would be expected to be capable to recruiting additional Aβ molecules into the β-sheet. The assembly pathway that produces the oligomer in our investigation is driven by interaction with the anionic detergent sodium dodecyl sulfate (SDS) near its critical micelle concentration. Initially 2-4 mer structures are formed. Removal of SDS, which is necessary for solid-state NMR measurements, results in an increase in aggregate size to 150 kDa (∼32-mers), as determined by multi-angle light scattering [16]. Additional experiments established that 150 kDa oligomers do not elongate in the presence of Aβ monomers to form amyloid fibrils, seed assembly of fibrils from soluble Aβ, or show enhanced fluorescence with thioflavin T (ThT) [17]. In contrast, protofibrils do undergo further assembly to fibrils and show enhanced fluorescence with ThT. The preparation and characteristics of 150 kDa oligomers share many features with those of globulomers [18] including their cell toxicity. For example, globulomers completely inhibited long-term potentiation in rat hippocampal slices and directly modulated recombinant P/Q-type and N-type calcium channels in HEK293 cells [19], and 150 kDa oligomers induced senescence in brain endothelial cells [20]. Furthermore, specific monoclonal antibodies raised against globulomers prevented synapse loss in a mouse model of AD [21]. We anticipate that knowledge of the 150 kDa oligomeric molecular structure will make it possible to understand the size limitation and to develop new diagnostic.

In this report, we present experimental evidence that there are two β-strands within the secondary structure of Aβ(1-42) when assembled into 150 kDa oligomers. The amino acid sequence for this peptide is DAEFRHDSGY^10^ EVHHQKLVFF^20^ AEDVGSNKGA^30^ IIGLMVGGVV^40^ IA. Previously, we reported that residues 30-42 form a β-strand (the C-strand) that is organized into an antiparallel β-sheet centered at residue V36 [22, 23]. New results in this report reveal the presence of a second β-strand spanning residues 11-24, which we call the N-strand. The antiparallel inter-molecular organization of the C-strand would motivate the hypothesis that the N-strand must also be arranged into an antiparallel β-sheet. However, NMR experiments designed to test this hypothesis, presented in this contribution, indicate otherwise.

## Results

### 2D ^13^C NMR spectra reveal two β-strands and evidence for multi-site occupancy

Spectral assignments, correspondences between ^13^C NMR peaks and isotopically labeled sites, were determined by collecting 2-dimensional (2D) ^13^C-^13^C NMR spectra with a short mixing time for dipolar recoupling on samples that were uniformly ^13^C-labeled (with ^13^C at every C site) within selected amino acids. For example, Figure 1 shows a 2D-fpRFDR spectrum [24] from a 150 kDa oligomer sample (Sample 1) that was ^13^C-labeled uniformly at K16, F20, V24, and G37. The pattern of off-diagonal peaks between directly bonded ^13^C atoms makes it possible to determine spectral assignments by analyzing crosspeak patterns that are unique to each uniformly labeled amino acid (see colored lines in Figure 1). Figures S1 and S2 show the 2D-fpRFDR spectra for samples prepared in this study, with labels chosen so that structure could be assessed for the whole peptide. Table S1 tabulates all ^13^C NMR peak positions (chemical shifts) and peak widths we have measured from spectra in Figures 1 and S1 (Samples 1-12). Table 1 reports the isotopic labeling employed for the full series of 150 kDa oligomer samples used in the study.

**Table 1.** Isotopic labeling employed for the 150 kDa oligomer samples analyzed in this study.

| **Sample** | **Isotopic Labeling** |
| --- | --- |
|  | **Uniform <sup>13</sup>C-labeling and <sup>15</sup>N- labeling at the indicated residues:** |
| 1 | K16, F20, V24, G37 |
| 2 | D7, G9, E11, L17, F19, A21 |
| 3 | E11, F19, I31, V36 |
| 4 | E11, L17, A21, M35, G38 |
| 5 | I32, M35, G37, V40 |
| 6 | Q15, V18, A21 |
| 7 | S8, Y10, V12, L34, G38, I41 |
| 8 | V12, E22, S26, N27, G33 |
| 9 | V12, F20, D23, K28, G29 |
| 10 | E11, H13, Q15, L17 |
| 11 | E11, K16, F19, V36 |
| 12 | A2, E3, F4, G9, V39 |
| 13 | F19, V24, G25, A30, I31, L34, M35 [22] |
|  | **Selective <sup>13</sup>C-labeling at the indicated sites:** |
| A | L17 CO |
| B | V18 CO |
| C | F19 CO |
| D | F20 CO |
| E | A21 CO |
| F | L17 CO and F19 CO |
| G | V18 CO and A21 CO |
| H | L17 CO and A21 CO |
| I | V18 CO and A21 CO (50% isotope-diluted) |

**Figure 1:**
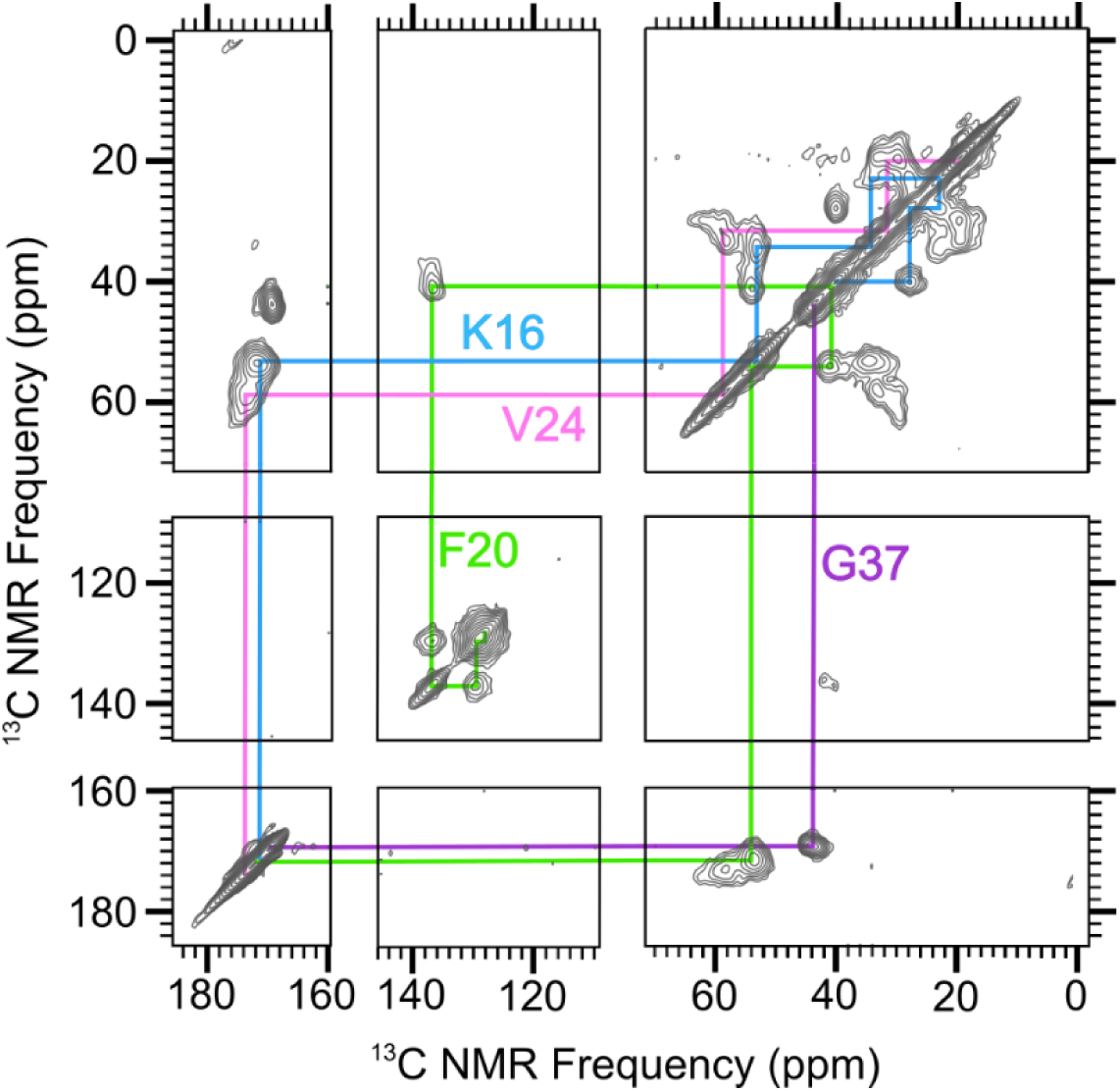
A 2D-fpRFDR spectrum of 150 kDa Aβ(1-42) oligomer sample that was uniformly labeled with ^13^C at K16, F20, V24, and G37. Colored lines indicate spectral assignments based on crosspeaks between directly bonded ^13^C atoms.

We assessed peptide secondary structure through analysis of ^13^C NMR frequencies (chemical shifts) for backbone C sites. Figure 2A reports secondary ^13^C chemical shifts for all CO, C^α^, and C^β^ for most of the residues in the oligomer and indicates the presence of 2 β-strand regions. Secondary chemical shifts are measured NMR peak frequencies relative to reported values for corresponding atoms in the same amino acids within model random-coil peptides in solution [25]. Secondary structure is known to correlate with ^13^C chemical shift when CO, C^α^, and C^β^ chemical shifts deviate in systematic ways from corresponding random-coil values for contiguous sequences of amino acids within the primary structure [26]. In particular, β-strand secondary structure typically observed in Aβ aggregates corresponds to negative CO and C^α^ secondary shifts and positive C^β^ secondary shifts. To be more concrete, we fed the assigned chemical shifts and peptide sequence to a computer program called TALOS-N [27] to predict the backbone torsion angles (ϕ/Ψ). For the 150 kDa oligomer, TALOS-N predicts the presence of two β-strands as shown in Figure 2A. The β-strands span residues 11-24 and 30-42; we refer to them as the N-strand and C-strand, respectively. The regions spanned by residues 1-10 and 25-29 are predicted to be an unstructured region and a turn, respectively. The detailed output of the TALOS program, which includes estimated uncertainty in torsion angle predictions, is shown in Table S2.

**Figure 2.**
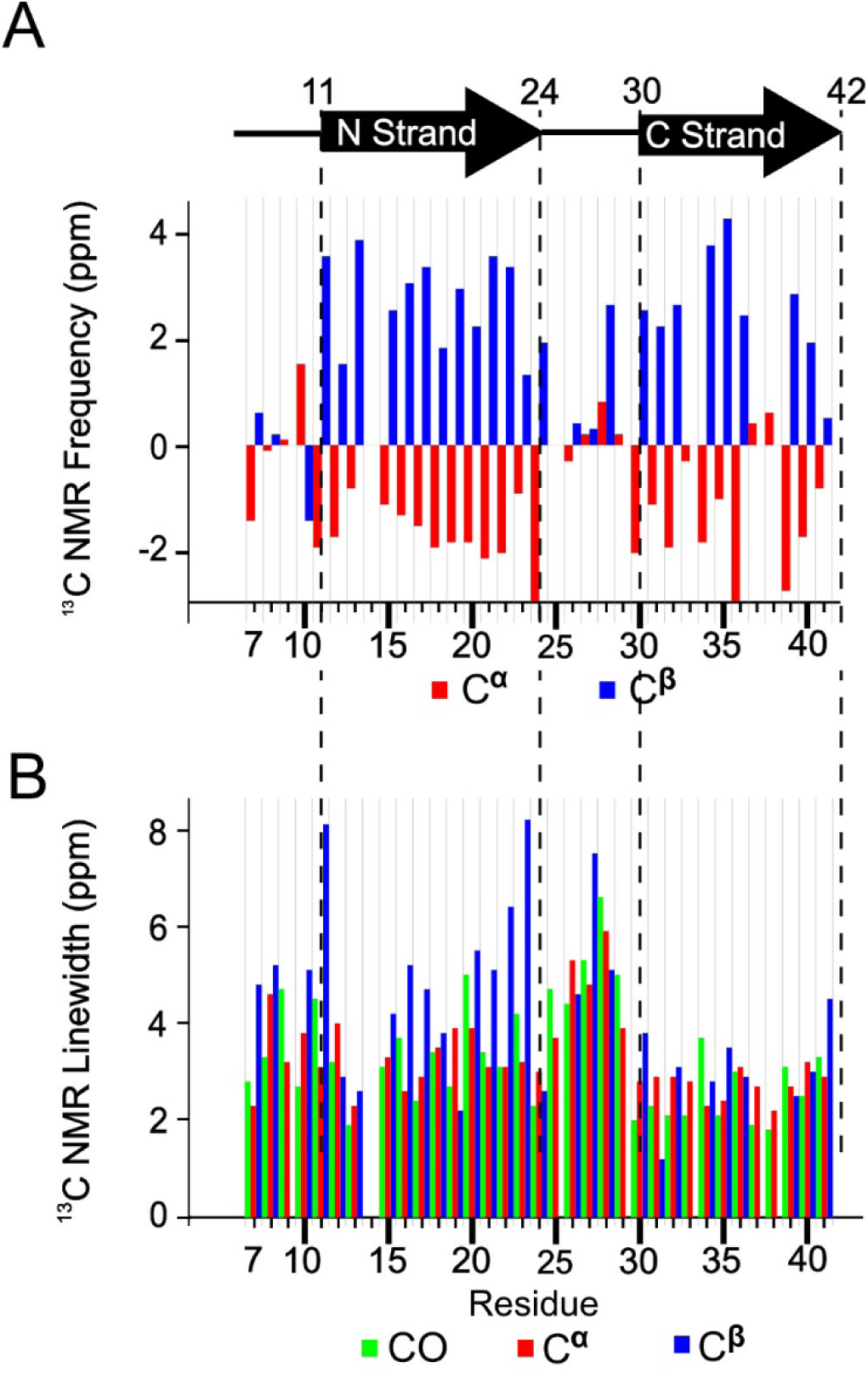
A) Secondary structure of peptide residues in the 150 kDa Aβ(1-42) oligomer, predicted by TALOS-N software, and the measured secondary ^13^C NMR backbone chemical shifts upon which the TALOS-N prediction is based. B) The NMR linewidths of CO (green), C^α^ (red), and C^β^ (blue) reported as full widths at half maximum measured by nonlinear regression of cross peaks in the 2D-fpRFDR ^13^C-^13^C NMR spectra to Gaussian functions.

An interesting observation is that the linewidths were larger for N-strand ^13^C signals than the C-strand counterparts, especially for the residues near the ends of the N-strand (Figure 2B). On average, the CO, C^α^, and C^β^ linewidths were 3.3 ± 0.5, 3.2 ± 0.5, and 4.0 ± 1.1 ppm (95% confidence region for full width half maximum), respectively, for the N-strand residues. The corresponding average linewidths for the C-strand were 2.5 ± 0.4, 2.7 ± 0.2, and 3.0 ± 0.7 ppm. Figure S3 shows that, based on a t-test, the CO and C^α^ linewidths differ significantly between the N- and C-strands. Figure S4 further illustrates this point, showing crosspeaks in 2D-fpRFDR spectra from the 6 valine residues in the peptide (3 within each strand). The NMR signals from the N-strand valine residues (V12, V18, and V24) exhibit more evidence of disorder via asymmetric 2D-NMR crosspeaks than the C-strand valine residues (V36, V39, and V40). In addition, although crosspeaks from the valines in N-strand have broader linewidths, the chemical shifts of the main Cα-Cβ crosspeaks of every valine are still consistent with β-strand conformation (Figure S4B). In terms of molecular structure, these spectra suggest that N-strand residues can occupy multiple magnetically inequivalent sites while C-strand residues do not. This inequivalence will be discussed further as more results are presented.

Most of our subsequent analysis of β-strand arrangements was based on the interpretation that N-strands and C-strands are arranged into distinct β-sheets in the oligomer. This assumption is supported by the aforementioned difference between the N- and C-strand ^13^C peak linewidths. Nevertheless, after presenting results that constrain relative orientations of adjacent N-strands, we do consider a β-sheet model in which N-strands and C-strands co-assemble into the same β-sheet.

### PITHIRDS-CT data and 2D-DARR negative results do not support in-register parallel or antiparallel N-strand β-sheet

^13^C-^13^C dipolar couplings obtained via the PITHIRDS-CT NMR experiment [28] (Figure 3A) indicate that N-strands are not assembled into an in-register parallel β-sheet. PITHIRDS-CT decays, or ^13^C NMR peak intensity as a function of the effective duration of ^13^C-^13^C dipolar recoupling, were measured for a series of oligomer samples that were each selectively ^13^C-labeled at a single backbone CO site within the N-strand (Samples A-E, Table 1). With only one ^13^C-labeled site per molecule, this measurement is expected to result in a decay curve that is bounded by the simulated (dashed) curves in Figure 3A if ^13^C atoms on adjacent molecules are separated by distances of approximately 0.5 nm. The measured decays are all considerably weaker than the simulated curves, indicating that we did not detect the influence of inter-molecular dipolar couplings. As detailed under Materials and Methods, the theoretical curves were obtained by simulating spin dynamics for 8 coupled ^13^C atoms arranged in straight lines with constant ^13^C-^13^C spacings of 0.5 or 0.6 nm. The theoretical curves were obtained by modeling the spin dynamics for systems of 8 ^13^C atoms under the influence of the PITHIRDS-CT pulse sequence; the ^13^C atoms were arranged into linear configurations and separated by a constant spacing of 0.5 or 0.6 nm. The linear geometry for ^13^C atoms was inspired by the in-register parallel β-sheet structure that is common to amyloid fibrils. In this configuration, a ^13^C atom placed at any backbone CO site or an alanine C^β^ site would be arranged into a linear configuration with a nearest-neighbor spacing of 0.48 nm. Discrepancies between PITHIRDS-CT experiments and simulations are known to occur [29, 30]. The reason(s) for the discrepancies are not fully understood, but possibly include deviations of atomic positions from idealized geometries, thermal fluctuations in positions of ^13^C atoms within the molecular ensemble, and spin relaxation for which the PITHIRDS-CT pulse sequence does not fully compensate. *Empirically, we have observed that a* ^*13*^*C PITHIRDS-CT curve that is bounded by the simulated curves indicates that each* ^*13*^*C atom in the sample is within approximately 0*.*5 nm from at least one other* ^*13*^*C atom*. Figure 3B demonstrates this empirical observation for previously published data on Aβ(1-42) amyloid fibrils [22], in which β-strands are known to form in-register parallel β-sheets [31]. The accompanying schematic illustrates the expected relative positions of ^13^C-labeled sites within the C-strands. We include more atomic details in model of the parallel β-sheet in Figure S5A.

**Figure 3.**
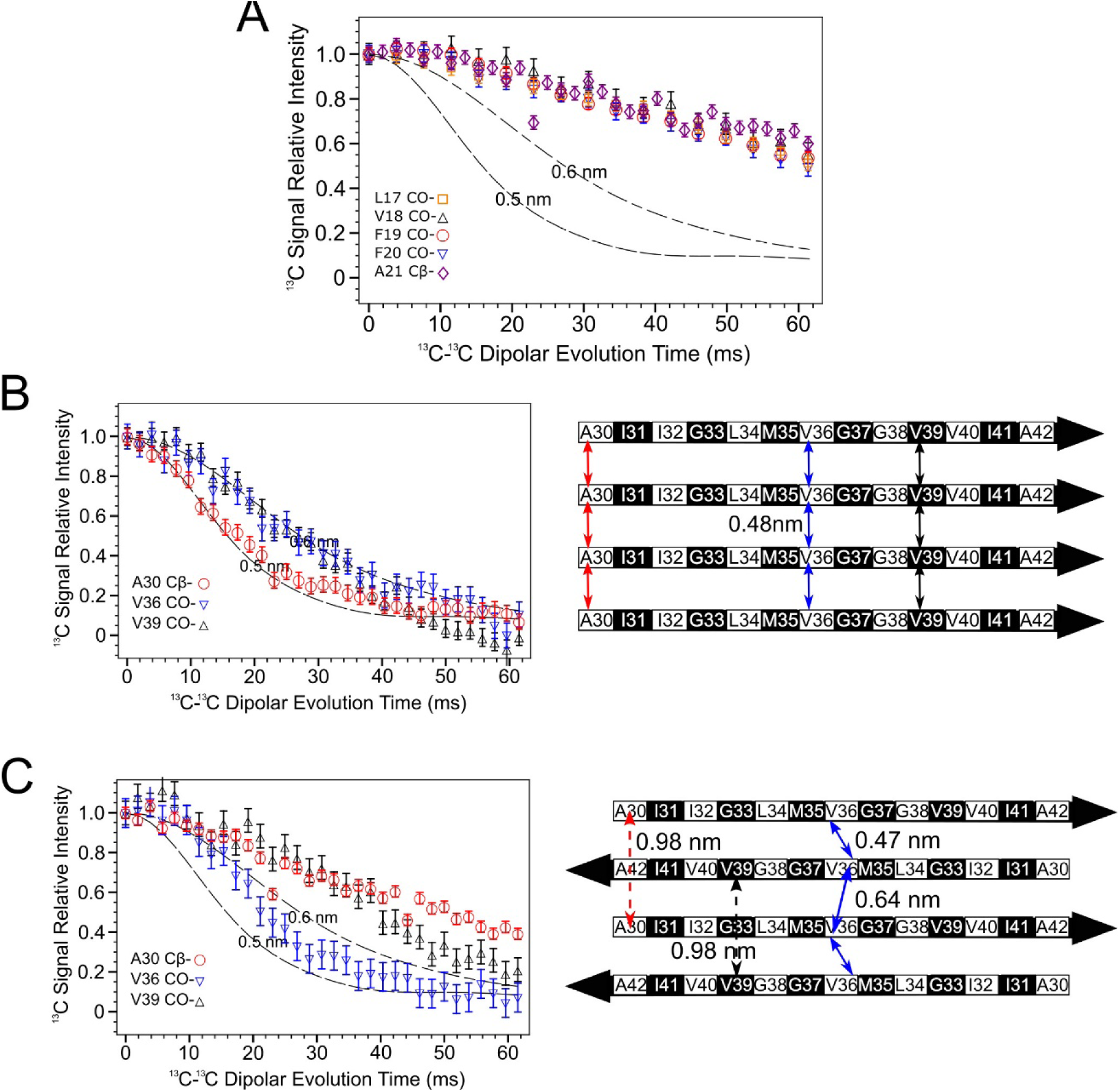
PITHIRDS-CT 13C-13C dipolar recoupling curves rule out an antiparallel alignment of N-strands. A) PITHIRDS-CT ^13^C-^13^C dipolar recoupling curves measured for 150 kDa Aβ(1-42) oligomer samples ^13^C-labeled at one backbone CO position per molecule within N-strand (Samples A-E). B) PITHIRDS-CT data from an Aβ(1-42) fibril (left) [22] and a schematic of an in-register parallel β-sheet formed by C-strand (right). The black and white coloring for each residue in the β-strand schematics indicate whether each sidechain is above (black) or below (white) the plane of the β-sheet. The double-headed colored arrows indicate that distances between equivalent backbone CO or C^β^ sites on adjacent molecules that are short enough (0.6 nm or less) to measurably affect PITHIRDS-CT decays. C) PITHIRDS-CT data on 150 kDa oligomer samples ^13^C-labeled at single near-backbone positions on C-strand (left) [22] and a schematic of the antiparallel C-strand β-sheet within 150 kDa oligomers (right). Since this β-sheet is centered at V36, the CO site of this residue would be the only backbone CO site that would yield a strong PITHIRDS-CT decay if selectively ^13^C-labeled (double headed arrows with solid line). Dashed lines in PITHIRDS-CT panels indicate simulated ^13^C interatomic distances that were calculated as outlined in the Methods. Black and white shading on the β-strand schematics indicate whether an amino acid sidechain is above or below the plane of the diagram, respectively.

The PITHIRDS-CT decays in Figure 3A are weak and exhibit no dependence on the residue position of the ^13^C-labeled backbone site, ruling out an antiparallel β-sheet for N-strands. The expected pattern of PITHIRDS-CT decays for an antiparallel β-sheet are illustrated with previously published data on the C-strands [22], which were determined to form an antiparallel β-sheet structure (Figure 3C; see also the all-atom β-sheet model in Figure S5A). Since V36 is at the center of the C-strand antiparallel β-sheet, ^13^C isotopic labeling of the CO site of V36 results in an expected nearest-neighbor ^13^C-^13^C distance of less than 0.5 nm and a PITHIRDS-CT decay that is similar to that observed for in-register parallel β-sheets. Although arrangement of ^13^C atoms in this case would not be linear, we expect the simulated curves to bound experimental decays observed for ^13^C spin systems in which nearest-neighbor ^13^C-^13^C distances are near 0.5 nm. For ^13^C-labeling at other backbone sites within the C-strand, the antiparallel configuration corresponds to significantly larger ^13^C-^13^C distances (above 0.7 nm). Correspondingly, weaker PITHIRDS-CT decays were observed with ^13^C labeling at the A30 C^β^ or the V39 CO site. Figure S6 illustrates that this behavior would be expected for any antiparallel β-sheet composed of β-strands of the same amino acids on different molecules: there is a single amino acid at the “center” with short inter-molecular distance between their backbone carbonyls. We also define each N-strand antiparallel β-sheet model by specifying the center residue and the sidechain orientations (Figure S6, S7 and S8). Based on the secondary structure determination for the 150 kDa oligomer (Figure 2A) and the knowledge that C-strands form an antiparallel β-sheet, we assumed the N-strands to also arrange into an antiparallel β-sheet with L17 or V18 at the center. However, the data in Figure 3A are consistent with no residue between L17 and A21 (inclusive) corresponding to the center of an antiparallel β-sheet.

In addition to PITHIRDS-CT experiments, we also used 2D-dipolar assisted rotational resonance (DARR) [32, 33] measurements to probe for a possible antiparallel N-strand β-sheet. This technique, when employed with a 500 ms mixing time for ^13^C-^13^C dipolar recoupling (500 ms 2D-DARR), produces spectra illustrated in Figure 4A and is capable of detecting 2D NMR crosspeaks corresponding to ^13^C atoms on distinct residues that are in dipolar “contact”. That is, we expected to observe inter-residue crosspeaks in 500 ms 2D-DARR spectra when *both residues are* ^*13*^*C uniformly labeled and at least one inter-residue pair of* ^*13*^*C atoms is within 0*.*6 nm* [34, 35]. We use contact charts like Figure 4B to facilitate a direct comparison between 2D-DARR NMR data and molecular modeling predictions. The predictions are based on a model of a Type-I antiparallel β-sheet centered at L17 (see Figures 4C, S6 and S7B). In the contact chart, each row and column correspond to the stretch of N-strand residues, and a grid square is shaded gray or red if the all-atom model predicts a corresponding inter-residue contact. Gray shading is for residue pairs that, by virtue of being close in sequence, can form intramolecular contacts (see Figure S9) and cannot be used to report on inter-molecular arrangements with the employed labeling schemes. In Figure 4A, the 2D-DARR contacts detected between L17 and F19 and between F19 and A21 provide examples of such intramolecular contacts and serve as positive controls. Red shading in the contact chart is for residue pairs that are predicted by the β-sheet molecular model to only form intermolecular contacts. Circles in Figure 4B and dashed arrows in Figure 4C indicate residue pairs which were uniformly ^13^C-labeled in the same sample but failed to produce detected inter-residue 2D-DARR crosspeaks. The spectrum in Figure 4A is marked to show where crosspeaks between E11 and A21, between E11 and F19, and between E11 and L17 would have been expected. More details of molecular modeling are shown in Figures S7 and S8; these figures describe a series of antiparallel β-sheet models centered at different residues within the N-strand. We show a contact chart for each model, indicating that we were unable to detect predicted 2D-DARR inter-residue contacts. The observed 2D-DARR negative results contradict every antiparallel β-sheet model with the center located between residues 14 and 18 (inclusive). Figure S10 exhibits selected slices taken from 2D-DARR spectra, with dashed rectangles indicating frequencies at which inter-residue crosspeaks were predicted but not observed. Although the absences of expected 2D-DARR crosspeaks may not be considered conclusive on their own (crosspeaks could be missing because of limited sensitivity), these negative results complement the PITHIRDS-CT data in Figure 3A as well as data showing observed inter-molecular 2D-DARR crosspeaks within the N-strand presented in the next section.

**Figure 4.**
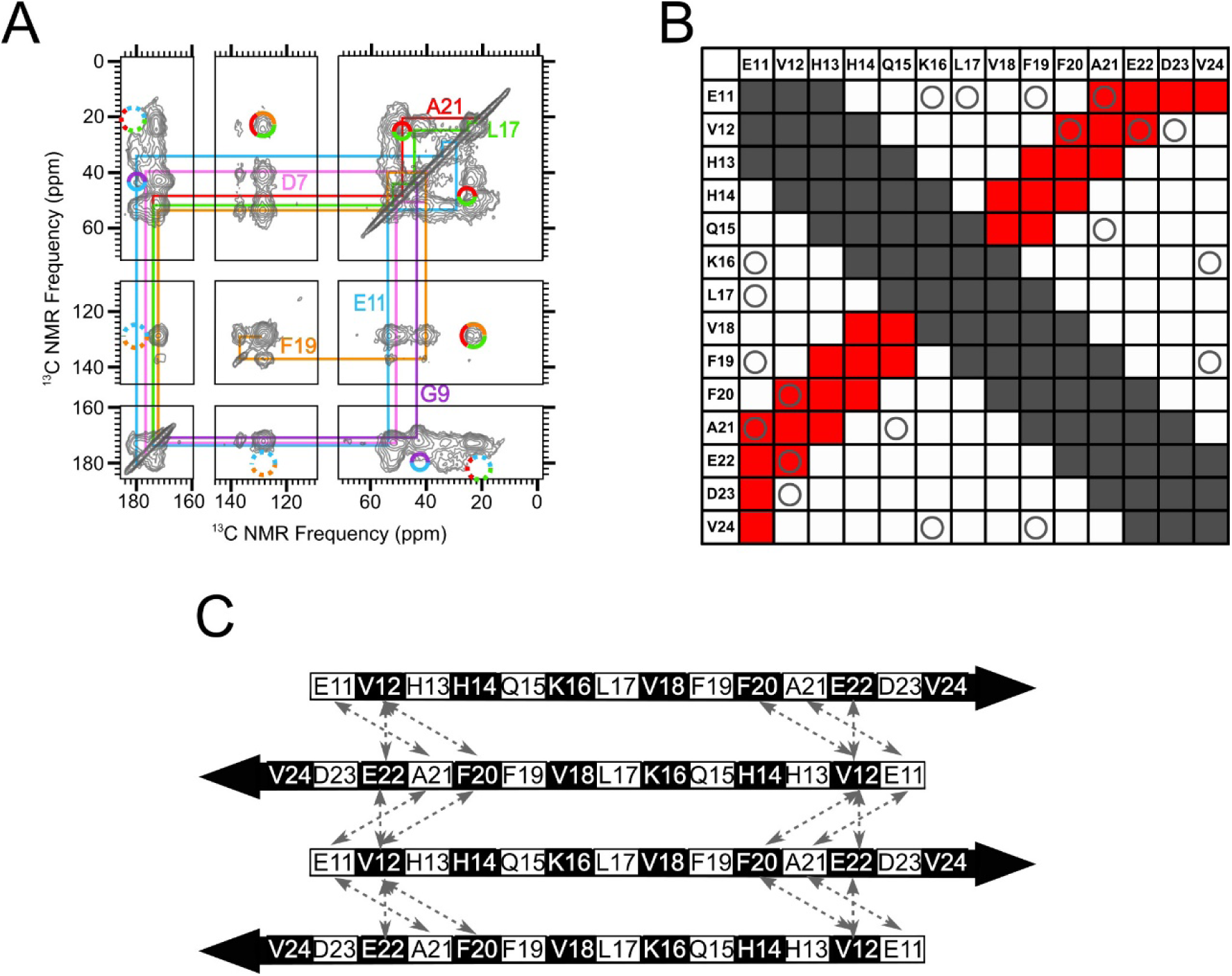
An in-register antiparallel β-sheet centered at L17 was not supported experimentally. A) A 500 ms 2D-DARR NMR spectrum of Sample 2, which was uniformly ^13^C-labeled at residues D7, G9, E11, L17, F19 and A21. To clarify spectral assignments, the colored lines indicate cross-peaks between directly bonded ^13^C atoms within each amino acid (intra-residue cross-peaks). The multi-colored circles drawn with solid lines indicate observed cross-peaks between ^13^C atoms on different labeled residues (inter-residue cross-peaks). The multicolored circles drawn with dotted lines indicate where inter-residue cross-peaks would be expected, based on an all-atom model of a N-strand antiparallel β-sheet centered at L17, but were not observed. (B) A contact chart that summarizes pairs of residues within the N-strand that are expected to exhibit inter-residue cross-peaks in 500 ms 2D-DARR spectra for an antiparallel β-sheet centered at L17. Squares colored gray indicate pairs with intra-residue cross-peaks or inter-residue cross-peaks that are uninformative because they occur through the primary sequence (atoms within a 0.5 nm distance). The squares colored red indicate pairs with predicted cross-peaks based on the model of an N-strand antiparallel β-sheet centered at L17. The “O” symbols represent the tested pairs from all ^13^C-labeled peptides that we examined which did not show cross-peaks. C) A schematic of an N-strand antiparallel β-sheet centered at L17. The dashed double-headed arrows indicate pairs of residues that correspond to circles in red squares in Panel B. Black and white shading on the β-strand schematic indicate whether an amino acid sidechain is above or below the plane of the diagram, respectively.

### 2D-DARR and PITHIRDS-CT results support out-of-register parallel β-sheet models for N-strand

Our inability to detect evidence for an antiparallel N-strand β-sheet compelled us to consider out-of-register parallel β-sheet models. Representations of all-atom parallel β-sheet models are shown in Figure S11 and S12. Schematic diagrams of these models will be used in the main text to guide the presentation of experimental results. We identify each parallel β-sheet model by the registry shift between adjacent strands: Models 2, 3, and 4 incorporate parallel N-strand β-sheets with registry shifts 2, 3, and 4, respectively. As will be discussed shortly, we also considered models in which the registry shift alternates in direction between adjacent molecules in the β-sheet. Parallel β-sheet models with alternating registry shifts will be designated as ±2, ±3, and ±4.

Figure 5A shows a 500 ms 2D-DARR spectrum of Sample 1. This sample was uniformly ^13^C-labeled at K16, F20, V24, and G37. With this spectrum, we observed inter-residue contacts between K16 and F20 and between F20 and V24. Since these contacts each correspond to a pair of residues that are 4 residues apart within the N-strand, they must arise from intermolecular ^13^C-^13^C dipolar couplings because intramolecular distances would be beyond the detectable range (see Figure S9). Additional 2D-DARR spectra showing detected inter-residue crosspeaks are presented in Figures S13-S16. A total of 9 inter-residue contacts, for N-strand residue pairs that are separated by 3 or 4 residues in sequence, were observed and shown as stars in Figure 5B. These positive results (namely, detected 2D-DARR contacts), as well as the negative results reported above (contacts anticipated but not observed), are consistent with the red shading in Figure 5B, predicted using Model 3 (Figure 5C and Figure S11B). Model ±3 (Figure 5D and Figure S12B) predicts a similar pattern of 2D-DARR contacts. Figures S11 and S12 further compare the pattern of observed and unobserved 2D-DARR contacts with Models 2, ±2, 4, and ±4; these models could also be consistent our 2D-DARR data. Thus, while our 2D-DARR data support the presence of registry-shifted parallel β-sheets, they do not precisely constrain the magnitude of the registry shift. It is also notable that the pattern of predicted 500 ms 2D-DARR contacts is not sensitive to alternation of registry shift direction. Nevertheless, there is an important difference between models with constant compared to alternating registry shifts: comparison of Figures 5C and 5D illustrates that alternation introduces magnetic inequivalence to the β-sheet structure. Taking F19 and its proximate residues for example, Figure 5C shows that, in Model 3, every F19 residue is adjacent to K16 on one neighboring strand and A21 on the other. In Model ±3 depicted in Figure 5D, half of the F19 residues are adjacent to two K16 residues on neighboring strands and half are adjacent to two E22 residues. Thus, an alternating registry shift would predict that isotopically labeled F19 residues would have two distinct sets of NMR peaks. Since we may lack the resolution to clearly distinguish between distinct sets of F19 peaks, this type of magnetic inequivalence could explain why we observed broader and more asymmetric NMR peaks for the N-strand than for the C-strand (see Figure 2B and Figure S4).

**Figure 5.**
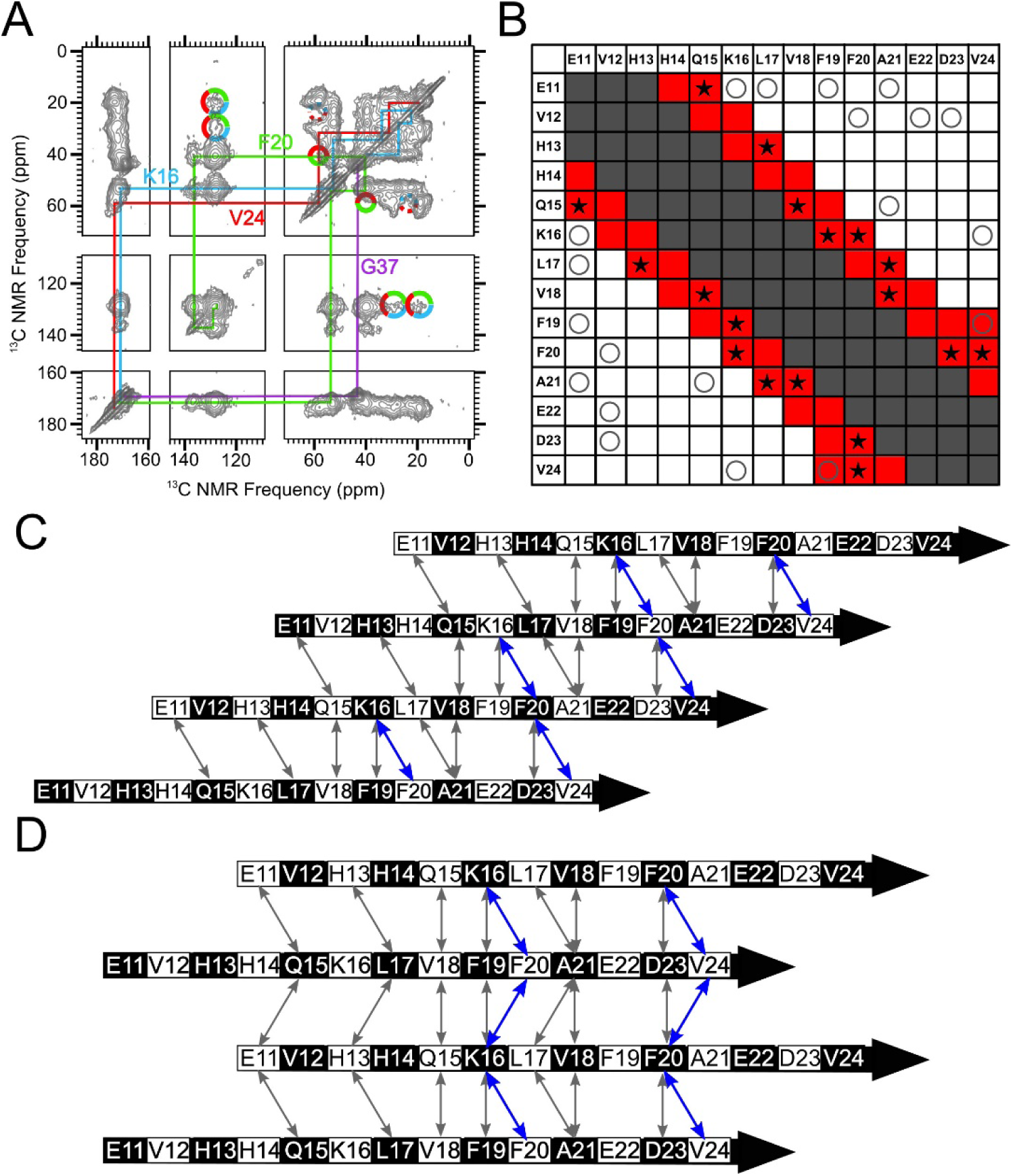
A parallel N-strand β-sheet shifted three residues out of register was consistent with experimental data. A) A 500 ms 2D-DARR spectrum of Sample 1, which was uniformly ^13^C-labeled at residues K16, F20, V24 and G37. The colored lines indicate intra-residue cross-peaks, and the multi-colored solid circles indicate observed inter-residue cross-peaks. The multi-colored dotted circles indicate where inter-residue cross-peaks may be anticipated, based on an all-atom model of a parallel N-strand β-sheet with +3 register, but were not observed. B) A contact chart similar to Figure 4B, but with red-colored squares corresponding to predicted 500 ms 2D-DARR contacts for Model 3. The star symbols (⋆) indicate test pairs whose cross-peaks were observed experimentally in 500 ms 2D-DARR spectra collected on samples in which the corresponding residues were both uniformly ^13^C-labeled. The “O” symbols and gray-shaded squares match their designations in Figure 4B. C) A schematic of Model 3. Black and white shading on the β-strand schematics indicate whether an amino acid sidechain is above or below the plane of the diagram, respectively. D) A schematic of Model ±3. The double-headed arrows in Panels C and D convey the same information as the star symbols in Panel B, and the residue pairs whose cross-peaks are observed in panel A (K16/F20, F20/V24) are highlighted in blue.

To validate our interpretation of the data in terms of the out-of-register parallel N-strand β-sheet models, we considered two alternative explanations of our experimental observations. The first possibility is that 2D-DARR experiments are more sensitive than we had expected, and the technique can detect crosspeaks between labeled residues that are separated by distances larger than 0.6 nm. If this alternative hypothesis were true, 2D-DARR crosspeaks detected between residues that are separated by 3 or 4 residues in sequence could be due to intramolecular ^13^C-^13^C dipolar couplings. To rule out this possibility, we considered a published 2D-DARR spectrum from an Aβ(1-40) fibril sample that was uniformly ^13^C-labeled at residues K16, F19, A21, E22, I32 and V36 [2]. It was shown previously that residues K16 and F19 in this structure are both within a β-strand that is organized into an in-register parallel β-sheet [2, 36]. We did not detect crosspeaks between K16 and F19 in the spectrum from this fibril, even though it exhibited a similar signal-to-noise ratio and narrower peaks compared to the spectrum in Figure 5A. For an in-register parallel β-sheet, the closest distance between K16 and F19 in the same strand would be 1 nm. This result supports the expectation that the 2D-DARR experiments would not detect crosspeaks between ^13^C-labeled residues separated by distances of 1 nm (3 residues) or longer. To further confirm the intermolecular nature of the K16/F19 contact observed in 150 kDa oligomers, we compared the 2D-DARR spectrum for oligomer Sample 11 (which contains uniform ^13^C-labels at residues E11, K16, F19, and V36) with that for Sample 11 diluted to 30% with unlabeled peptide (Figure S17). This comparison showed attenuation of the crosspeaks between K16 and F19 as a result of isotopic dilution, indicating that the crosspeaks between K16 and F19 are due to ^13^C-^13^C dipolar couplings between labeled residues on different molecules.

The second alternative explanation is that that the observed inter-residue 2D-DARR crosspeaks are due to stacking of distinct N-strand β-sheets rather than contacts between neighboring β-strands in the same β-sheet. To examine this explanation and obtain more specific constraints on N-strand backbone alignment within the β-sheet, we performed PITHIRDS-CT measurements on samples that were each selectively ^13^C-labeled at two backbone CO positions in the N-strand. With ^13^C-labeling only at backbone sites, PITHIRDS-CT results report specifically on organization of adjacent β-strands within the same β-sheet: peptide backbone atoms on different stacked β-sheets would be separated by distances that are too large (> 0.7 nm) to affect PITHIRDS-CT decays. We previously employed a similar strategy to detect out-of-register parallel β-sheet structure within nanofibers of the RADA16-I designer peptide (amino acid sequence RADA^4^RADA^8^RADA^12^RADA^16^ with acetylated and amidated N- and C-termini, respectively) [37]. ^13^C-Labeling at a single A8 CO site on the β-strand backbone of RADA16-I fibrils yielded a weak PITHIRDS-CT decay that is not consistent with an in-register parallel β-sheet. With a different RADA16-I sample that was labeled with ^13^C at two CO sites separated by 2 residues (A4 and A6), we were able to detect a PITHIRDS-CT decay that is consistent with a parallel β-sheet and a registry shift of 2 (Figure S18A and B). A similar PITHIRDS-CT decay analysis was conducted on the series of 150 kDa oligomer Samples F, G, and H that were ^13^C-labeled at two CO sites within the N-strand (Table 1). In comparison to the decay observed with ^13^C-labeling of the oligomer at only one site (Sample A), samples with pairs of ^13^C-labeled CO sites exhibited measurably stronger PITHIRDS-CT decays (Figure 6A). For Samples F, G, and H, the ^13^C-labeled sites were either 2 residues apart (Sample F: L17 and F19), 3 residues apart (Sample G: V18 and A21), or 4 residues apart (Sample H: L17 and A21). Stronger PITHIRDS-CT for Samples F, G, and H relative to Sample A supports the presence of an out-of-register parallel N-strand β-sheet. The registry shifts introduce intermolecular ^13^C-^13^C couplings. It should be noted that the PITHIRDS-CT curve for labeling of RADA16-I at only a single CO site (A8) per molecule (Figure S18A) resulted in a stronger PITHIRDS-CT than the analogous two CO-site labels applied to Aβ(1-42) 150 kDa oligomers. Unlike the oligomers, RADA16-I forms a single β-strand per molecule, and β-sheets stack along compact interfaces composed only of alanine methyl sidechains. Consequently, the distance between A8 CO sites on different β-sheets is in the detectable range (0.63 nm) and shorter than would be possible with oligomers. Further analysis, presented in the next section, is necessary to explain why observed decays are weaker than the theoretical curves.

**Figure 6.**
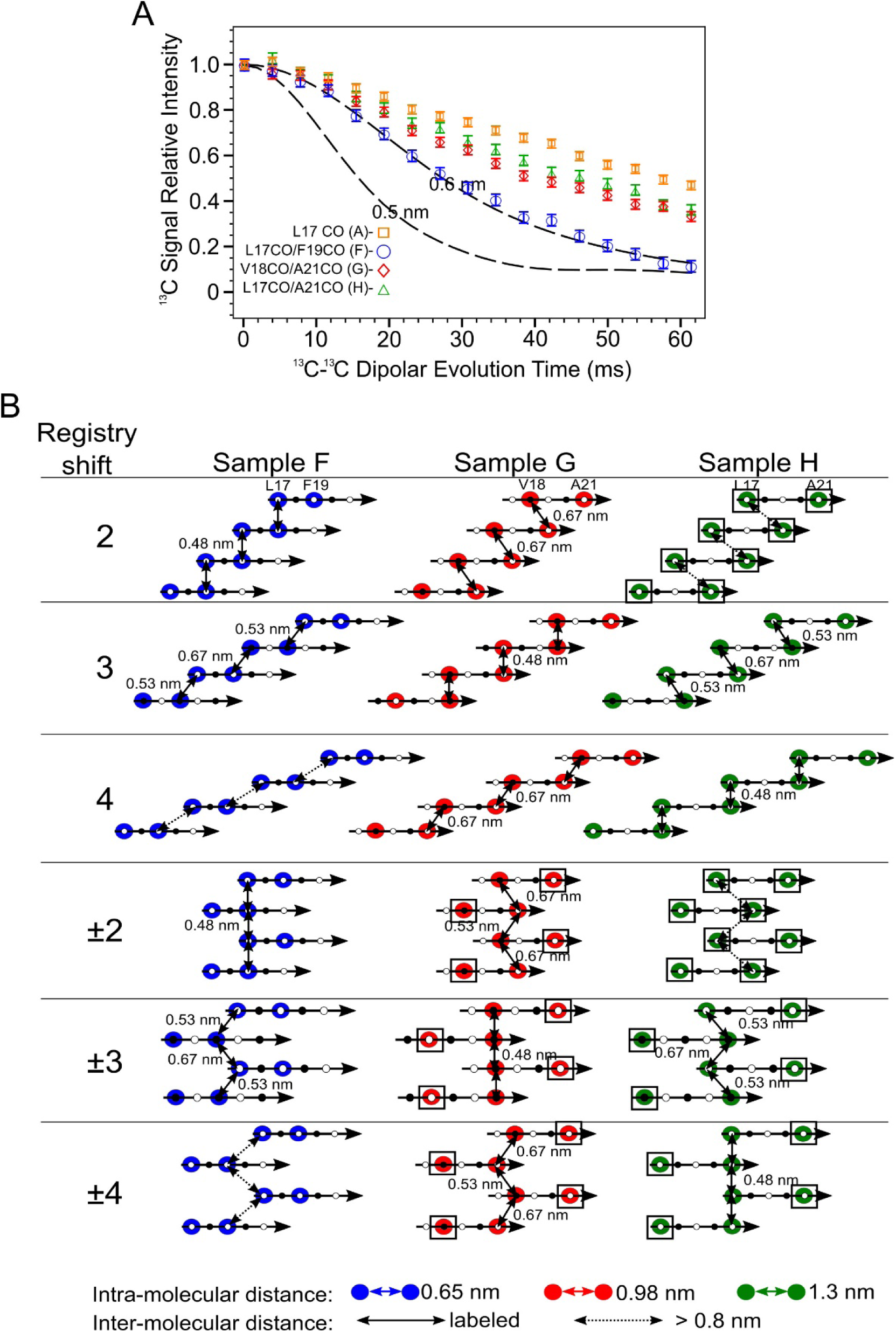
Doubly ^13^CO labeled PITHIRDS-CT data indicates out-of-register alignments of the N-strands. A) PITHIRDS-CT data for 150 kDa oligomers ^13^C-labeled at two backbone CO positions within the N-strand (Samples F, G and H). For comparison, the PITHIRDS-CT curves for Sample A (^13^C at L17 CO) is also plotted. Dashed lines in Panels A and C were the same simulated curves as in Figure 3. B) Diagrams illustrating the relative positions of ^13^C-labeled CO sites for Samples F, G and H, predicted by Models 2, 3, 4, ±2, ±3 and ±4. Colored circles indicate residues in which CO sites are ^13^C-labeled. Doubled headed arrows indicate ^13^C-^13^C distances between the labeled sites. Boxes around circles indicate positions of uncoupled spins.

### Registry shift within the N-strand β-sheet is likely to be 3 residues and alternate in direction (±3)

The PITHIRDS-CT decays in Figure 6A can be rationalized in terms of relative atomic positions predicted by molecular models of out-of-register parallel β-sheets. Figure 6B illustrates the relative positions of the ^13^C-labeled sites for Samples F, G, and H within candidate models with registry-shifted parallel N-strand β-sheets. More detailed depictions, including representations of all-atom β-sheet models, are shown in Figure S11 and S12. Figure 6B also shows how relative atomic positions change between models with different registry shifts.

Two limiting cases for PITHIRDS-CT decays illustrate the incompatibility of measured PITHIRDS-CT decays with most of the registry-shifted parallel N-strand β-sheet models considered here. The simulated curves in Figure 6A correspond to one limiting case: these curves bracket the expected PITHIRDS-CT decays for systems in which every ^13^C-labeled site experience a dipolar coupling with at least one other ^13^C-labeled site within 0.48 nm. The result would be a “fully coupled” ^13^C spin system of isotopically labeled sites with the strongest possible ^13^C-^13^C couplings for the structures being considered. The single ^13^C-labeled Aβ(1-42) fibrils (Figure 3B) and the doubly ^13^C-labeled RADA16-I nanofiber (Figure S17B) are two examples of fully coupled spin systems. At the other extreme, we call a ^13^C spin system “fully uncoupled” if every ^13^C-labeled site is at least 0.7 nm from every other ^13^C-labeled site. For a fully uncoupled ^13^C-spin system, we would expect to observe a PITHIRDS-CT decay that is indistinguishable from the decay observed with only one backbone CO ^13^C site per molecule (Samples A-E, Figure 3A). Even when a spin system is not fully uncoupled, we refer to ^13^C atoms as “uncoupled spins” when they are at least 0.7 nm away from all other ^13^C-labeled sites (marked by black boxes in Figure 6B). Note that fully uncoupled ^13^C spin systems do exhibit measurable PITHIRDS-CT decays, most likely due to nuclear spin relaxation effects and dipolar couplings to the 1% natural abundance ^13^C background on adjacent C sites. *The PITHIRDS-CT decays measured for Samples F, G, and H are inconsistent with both the fully coupled and fully uncoupled extreme cases*. The decays for these samples are all weaker than the theoretical curves that correspond to fully coupled spin systems, but they are measurably stronger than the decays measured for samples ^13^C-labeled at one backbone CO site per molecule. In addition, we verified the presence of inter-molecular ^13^C-^13^C dipolar couplings in Sample G is inter-molecular with a PITHIRDS-CT experiment on an isotopically diluted sample (Figure S19). Models 2, 3, and 4 each predict a fully coupled ^13^C spin system for one sample among Samples F, G, and H, while Models 2 and ±2 predict the fully uncoupled ^13^C spin systems for Sample H (Figure 6B). Since these predictions of spin systems are not confirmed in experiments, only two registry-shifted parallel β-sheet models can be consistent with the data because they predict PITHIRDS-CT decays for Samples F, G, and H that are in between the limiting cases; these are Models ±3 and ±4.

Figure 7 compares measured PITHIRDS-CT decays from Samples F, G, and H to simulated curves that are revised using ^13^C atom coordinates in Models ±3 and ±4. This comparison, along with trends predicted by relative atomic positions (Figure 6B), points to Model ±3 as the most likely model to fit the data. The clearest distinction between Model ±3 and Model ±4 is given by the prediction of the PITHIRDS-CT decay for Sample F. Model ±3 predicts a curve for this sample that more closely resembles the experimental curve. Figure S20 shows that the Sample F PITHIRDS-CT decay predicted using Model ±4 is well-represented by the decay of a pair of ^13^C atoms separated by the intra-molecular ^13^C-^13^C distance of about 0.65 nm because this model predicts a longer minimum inter-molecular ^13^C-^13^C distance of 0.84 nm. The observation that the measured Sample F PITHIRDS-CT decay is stronger than the prediction based on Model ±4 indicates the influence of inter-molecular ^13^C-^13^C dipolar couplings, as predicted by Model ±3. Models ±3 and ±4 predict similar PITHIRDS-CT decays for Sample G (^13^C-labeled backbone CO sites 3 residues apart) and Sample H (^13^C-labeled sites 4 residues apart), with both models predicting that 50% of the ^13^C spins are uncoupled. Model ±3 does predict slightly shorter ^13^C-^13^C distances for Sample G than for Sample H, but the difference between these two measured curves is not large enough to overcome the uncertainty in the analysis.

**Figure 7.**
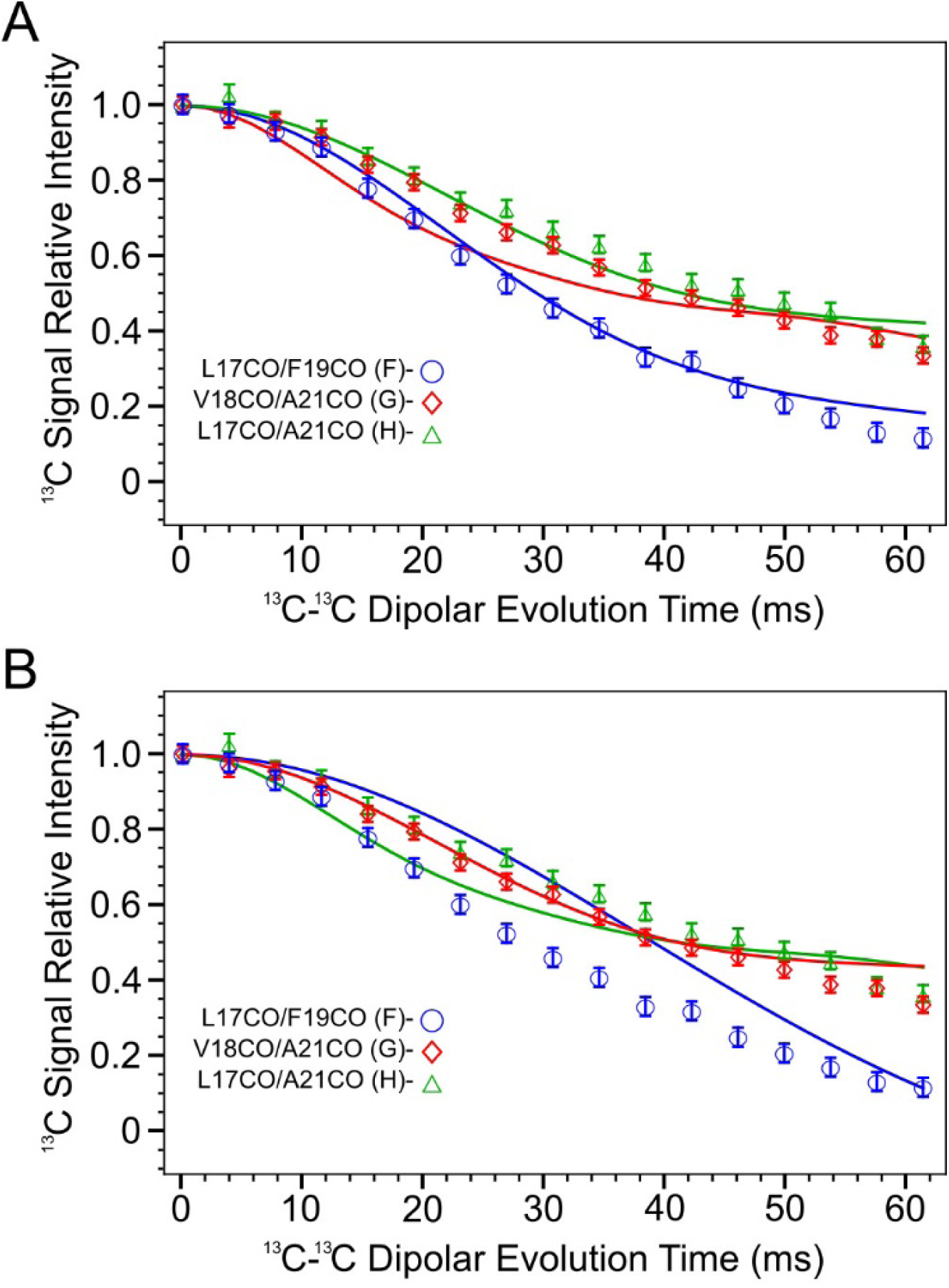
Simulations of PITHIRDS-CT curves for samples F, G and H according to ^13^CO atom coordinates in model ±3 and ±4. The simulated curve has the same color as the corresponding data series (blue: sample F, red: sample G, and green: sample H). A) Model ±3. B) Model ±4.

## Discussion

The most significant result of the present analysis is that the 150 kDa Aβ(1-42) oligomer structure includes both antiparallel and parallel organization of neighboring β-strands. Our previously published work reported 2D NMR contacts and ^13^C-^13^C dipolar recoupling data that are consistent with antiparallel arrangement of neighboring C-strands centered at V36. In the present study, similar experimental techniques applied to the N-strand did not reveal the expected antiparallel N-strand arrangements; inter-residue 2D-DARR contacts anticipated by this arrangement were not observed. Instead, we detected inter-molecular 2D-DARR contacts between N-strand residues that are consistent with a registry-shifted parallel β-sheet. If one assumes that β-sheets containing N-strands include no C-strands, Figure 8A is the N-strand β-sheet model that agrees best with the 2D NMR and ^13^C-^13^C PITHIRDS-CT data presented thus far. To summarize the evidence for this figure, the β-strand secondary structure spanning residues 11-24 is based on TALOS-N predictions from CO, C^α^, and C^β^ NMR ^13^C chemical shifts (Figure 2). The registry shift of 3 amino acids between adjacent strands is our interpretation of the pattern of 500 ms 2D-DARR inter-residue contacts between residues within the N-strand (Figure 5) and the weakly coupled PITHIRDS-CT decays for Samples F, G, and H (Figure 6). The alternating pattern of registry shifts (±3) yields the best agreement between experimentally measured decays of ^13^C NMR signal intensity of PITHRIDS-CT experiments on samples that were ^13^C-labeled at two backbone carbonyl sites within the N-strand (Figure 7).

**Figure 8.**
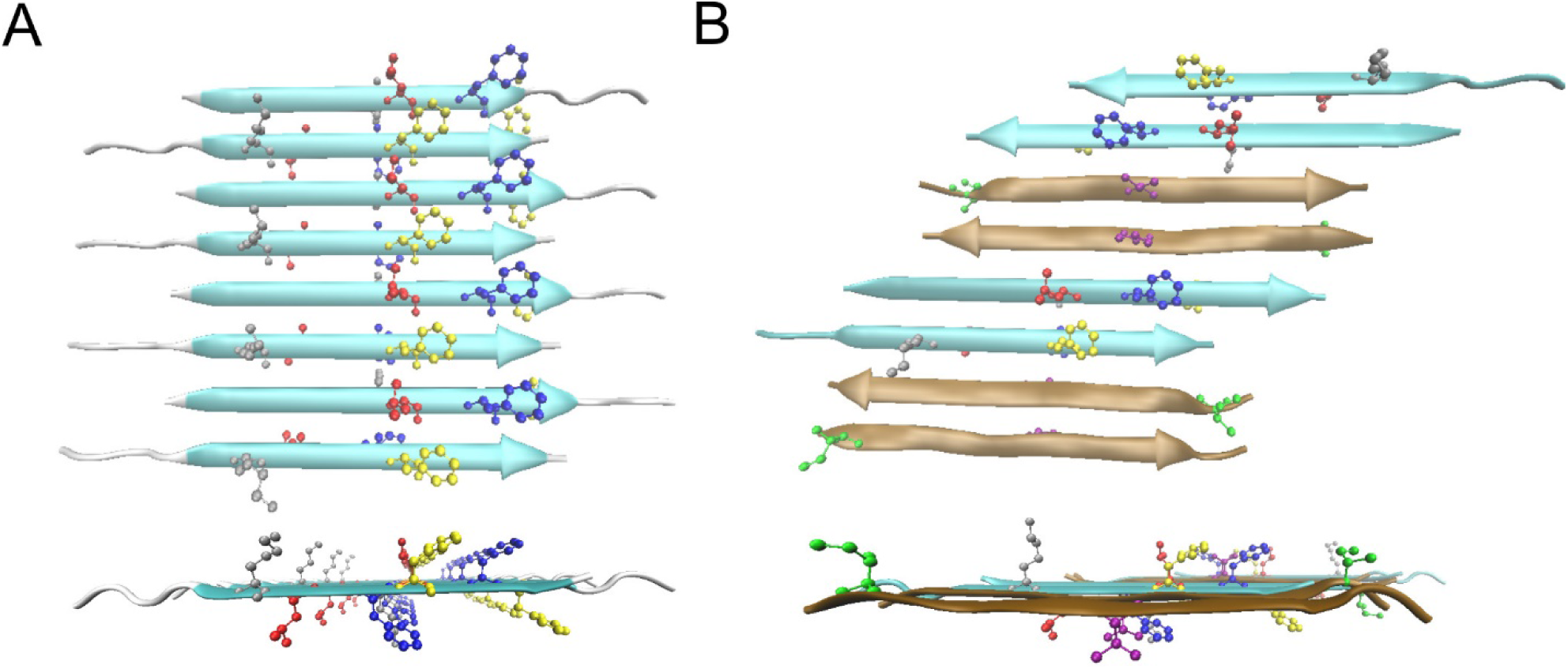
Possible β-sheet structures in the Aβ(1-42) 150kDa oligomers. A) The top and side views of the all-atom structural model of Model ±3 of N-strands. The color code for residues are: Grey K16, Red L17, Blue F19, Yellow F20. B) The top and side view of one structural model of a mixed β-sheet formed by both N-strands and C-stra nds. The N-strands are colored in cyan and the C-strands are in ochre. The color code for residues are: Grey K16, Red L17, Blue F19, Yellow F20, Green I31, Purple V36.

An alternative model that could explain many of the experimental observations in this study is shown in Figure 8B. In this model, each β-sheet contains both N-strand and C-strands. Such a configuration could result from Aβ(1-42) molecules in β-hairpin conformations, as has been proposed by Hoyer et. al. [38] and Doi *et. al*. [39], or from domain swapping as has been proposed by Stroud *et. al*. [40]. We did not consider this structure in earlier sections because we interpreted the broader ^13^C line widths for the N-strand compared to the C-strand to mean that the two β-strands were organized into different β-sheets. However, the model in Figure 8B does retain adjacent pairs of C-strands organized antiparallel and pairs of N-strands organized in parallel and out-of-register as shown in Figure 8A. Further examination of the model in Figure 8B, given in Figure S21 and S22, indicates that this model would predict the observed 500 ms 2D-DARR contacts (Figure 5B) and the ^13^C-^13^C PITHIRDS-CT data (Figures 3A and 6A) between residues within the N-strand and remain consistent with our previously published work that specified antiparallel organization of adjacent C-strands in the structure [23]. The model in Figure 8B also predicts some inter-residue crosspeaks in 500 ms 2D-DARR spectra between residues in N- and C-strands. Spectra from the labeled 150 kDa oligomer samples in Table 1 were examined for such crosspeaks, and inter-residue contacts were detected for 10 pairs and absent for 5 other pairs (Figure 9). If one particular registry between N- and C-strands is analyzed, *e*.*g*., one with the observed close contact between F19 in the N-strand and V36 in the C-strand, the contact chart between N- and C-strands in Figure S21C can be generated. This chart shows that one other observed contact between L17 and G38 is predicted, while 8 others are not. Therefore, models like that in Figure 8B are incomplete because we have yet to rationalize the pattern of 2D-DARR contacts between N-strand and C-strand residues in Figure 9. We suspect that constraints on oligomer nanoscale dimensions are necessary in order to fully understand the oligomer molecular structure. We will subsequently compare these structural constraints with previously published structural data on Aβ fibril, protofibrils, and oligomers. However, we are aware of no previously proposed molecular structural model that simultaneously includes both parallel and antiparallel intermolecular alignments of β-strands.

**Figure 9.**
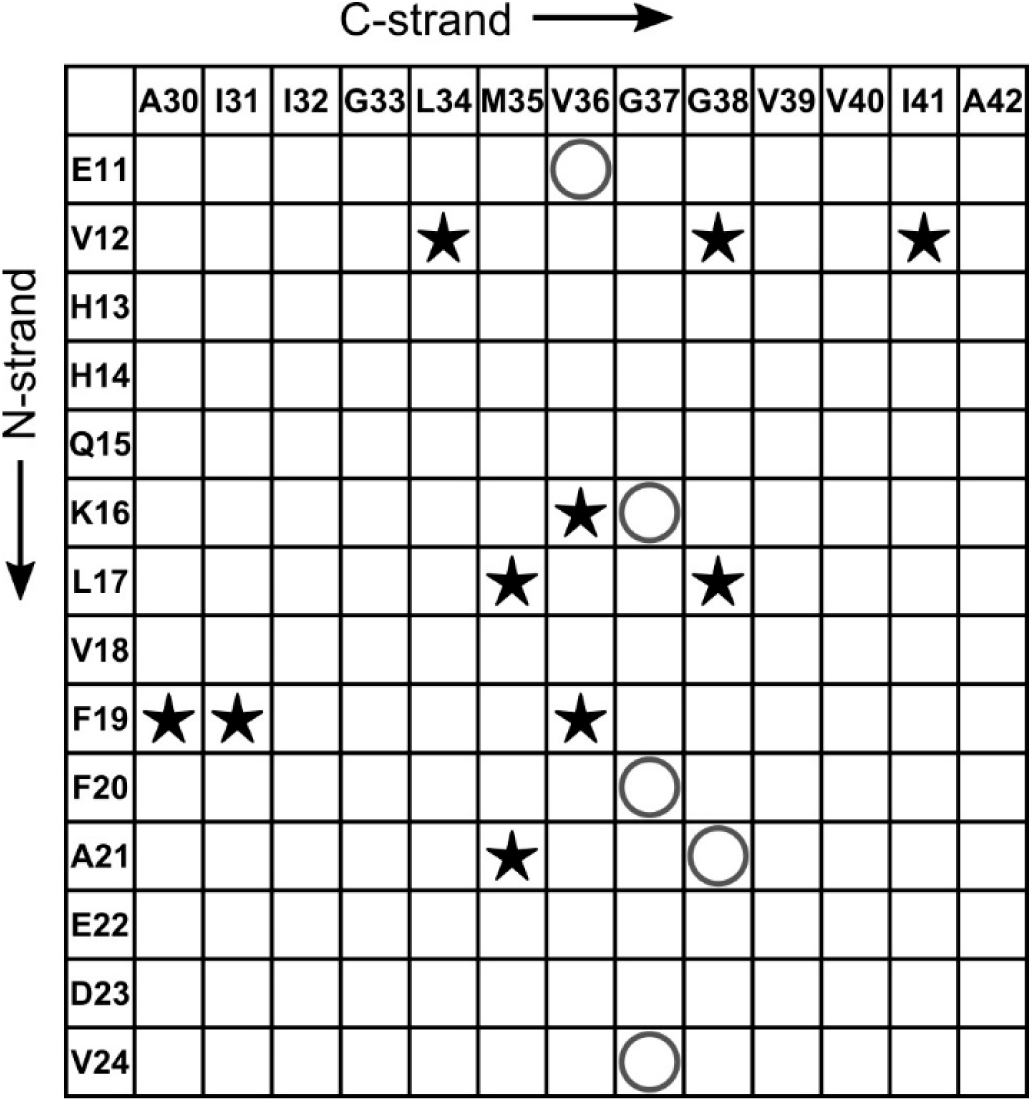
A contact chart that summarizes the experimentally detected and non-detected interactions between the residues in N-strand and that in C-strand. The star symbols (⋆) indicate pairs of residues whose cross-peaks were observed in 500 ms 2D-DARR spectra, while the “O” symbols represent the pairs from all ^13^C-labeled peptides that did not show cross-peaks in spectra.

The best-studied Aβ aggregate structures are amyloid fibrils – the main component in amyloid plaques. These are nanofibers each composed of thousands of Aβ molecules, with fibril widths between 5 and 10 nm and lengths ranging from 100 nm to above 1 μm [41]. In the literature, there is some detailed molecular structures of an Aβ(1-42) amyloid fibril [5-9] and a number of Aβ(1-40) fibril structural models [2, 42] based on well-defined experimental constraints on β-strand secondary structure and inter-strand organization. In each published model, every molecule adopts the same conformation within the amyloid fibril, with 2 or 3 β-strand domains per molecule arranging into “U-shaped” or “S-shaped” conformations (Figure S23A and B). In these fibril structures, β-strands organize into β-sheets such that each β-sheet is composed of equivalent β-strand segments (e.g., a β-strand formed by residues 12 to 23). The result is that β-sheets are close to planar (some twist can be observed along the fibril) and can extend to include thousands of β-strands. Most amyloid fibrils are composed of in-register parallel β-sheets, but Qiang *et. al*. [10] showed that the Iowa mutant (D23N) of Aβ(1-40) can form a fibril composed of either antiparallel β-sheets or in-register parallel β-sheets depending on the assembly conditions (Figure S23C). It is notable that the planar nature of an amyloid fibril appears to necessitate either that all β-sheets in a fibril be composed of parallel β-strands or that all β-sheets be composed of antiparallel β-strands. We are aware of no instance of co-existence of parallel and antiparallel β-sheets in one fibril structure. The presence of parallel and anti-parallel β-sheet in one structure, just as in our 150kDa Aβ oligomers, could possibly suppress amyloid fibril development.

Other conditions of Aβ aggregation generate protofibrils, which can undergo further assembly to fibrils and show enhanced fluorescence with ThT. The few NMR structural studies that have been conducted on Aβ protofibrils have not been very informative, but some of them have proposed that the protofibrils contain a β-hairpin structure (an intra-molecular antiparallel β-strand) [39, 43] and that they would convert into mature amyloid fibrils by conformation changes. A more recent NMR study revealed a hexameric barrel as the building block of a protofibril sample [44] with an engineered disulfide-containing Aβ(1-42) that locks into a peptide β-hairpin (Aβcc) [45, 46]. These hexameric barrels can interact to form elongated protofibrils that resemble wild type Aβ(1-42) protofibrils, but they cannot proceed to fibrils. Proposed β-hairpin secondary structures that are not locked by covalent crosslinks and their hydrophobic interactions in protofibrils could in principle rearrange to the configuration of mature fibrils, but there is as yet no experimental evidence of this. β-Hairpins also could in principle arrange parallel and anti-parallel β-strands into mixed β-sheets with some similarity to those in Figure 8B, and we don’t have enough experimental data to completely exclude all such structures in 150 kDa oligomers. However, such β-hairpins may not be building blocks that are stable enough to resist conversion to fibrils.

The “Aβ oligomer” is the general term for smaller Aβ aggregates, and it involves a wide range of sizes from dimers to protofibrils. Conformation-selective antibodies distinguished two types of endogenous oligomers [47] and surveyed their distribution in the brain. Both types have analogues among synthetic oligomers produced *in vitro*, but the synthetic oligomers can be produced in larger amounts that allow their structural characterization at the molecular level. One type is closely related to the amyloid fibril. This type may act as intermediates in the fibril aggregation process [48] or be the product of fibril-induced Aβ assembly [49]. Studies on some oligomers of this “intermediate” type found they usually have in-register parallel β-sheets like fibrils [47, 48]. Tycko and co-workers [50] used solid-state NMR to show that the fibril-like “U-shape” conformation of the Aβ peptide was already formed in the early oligomer stage. Based on solid-state NMR data, Tycko has suggested that early Aβ oligomers contain antiparallel β-sheets that are “off-pathway” for fibril formation and that these oligomers dissociate before reaggregating as “on-pathway” oligomers with parallel β-sheet structure [51]. Off-pathway oligomers correspond to the second type of oligomer noted above, with assembly pathways and secondary structures different from those of amyloid fibrils, making them “fibril-irrelevant”. Aggregation conditions play a significant role in determining whether this type of Aβ oligomer can be detected *in vitro*. Initial treatment of synthetic Aβ(1-40) or Aβ(1-42) at high pH and monomer isolation by a technique like size exclusion chromatography (SEC) are necessary to insure that residual aggregates are eliminated [52]. Subsequent aggregation in dilute salt buffers like buffered saline lead to fibril formation through protofibril intermediates [53]. No other oligomers are detected by biophysical methods during such aggregation unless sensitive techniques like those involving Microfluidic modulation spectroscopy (MMS) to detect minor components [54]. To obtain *in vitro* preparations that are largely off-pathway oligomers, it is necessary to treat monomeric Aβ(1-42) with agents that promote aggregation. One such agent is DMSO (footnote^1^), which generates oligomers called Aβ-derived diffusible ligands (ADDLs) [56]. A second is SDS at a concentration just below its critical micelle concentration, which leads to the 150 kDa oligomers described here [17]. It is noteworthy that only Aβ(1-42) and not Aβ(1-40) forms these off-pathway oligomers [17, 18, 57]. ADDL preparations contain heterogenous Aβ(1-42) off-pathway oligomers and display two SEC peaks corresponding to large aggregates (65-80 kDa) and monomers [12, 58]. No ADDL characterizations by NMR have been reported.

The 150kDa Aβ oligomers are no doubt fibril-irrelevant oligomers (or off-pathway for fibril formation). First, the oligomer samples are very stable and cannot spontaneously convert into fibrils even in the presence of Aβ(1-42) monomers. Second, Aβ(1-42) peptide molecules first form dimers to tetramers during preparation and then further assemble into 150 kDa oligomers [17]. The final oligomer sample shows little fluorescence response to ThT, which indicates no fibril is involved in the formation of the oligomer. Third, the secondary structures deduced from NMR in our oligomer samples are fundamentally distinct from those in the fibrils. For metastable Aβ oligomers in protein-like size (50∼200 kDa), the 150 kDa structure may demand more compact folding rather than growth capability, and thus the difference in secondary structure is expected.

Remarkably, the out-of-register parallel β-sheet structure, with alternating registry shifts of +3 and -3 in the N-strand region, that we deduce for 150 kDa oligomers is a very unusual arrangement. Although there are few examples of structures with such shifts, one is the out-of-register anti-parallel β-sheets formed by Aβ11– 25 fibrils formed at pH 2.4 [59]. The β-strands in these fibrils correspond to the N-strand region in 150 kDa oligomers but are anti-parallel rather than parallel. A second example is the model peptide ccβ-p, which has pH-dependent registry shift numbers in anti-parallel β-sheets that form fibrils [60]. One registry shift is +3, and this odd number registry shift would create a flip-over between neighboring β-strands. The flip-over would require that sidechains from the same residue on adjacent molecules alternately point up and down within one β-sheet (see Figure S12). The N-strand region of Aβ(1-42) and the ccβ-p model peptide (Ac-SIREL EARIR ELELR IG-NH2) both contain several charged sidechains, and thus the pH dependence indicates that sidechain charges may motivate the registry shifts. Further tests are needed to confirm this assumption. It is also worth noting that secondary involving antiparallel and out-of-register parallel β-sheets have been reported in other oligomers. Raussens and co-workers [61] observed a conversion of anti-parallel β-sheets in Aβ(1-42) oligomers to parallel β-sheets in Aβ(1-42) fibrils by FTIR. Eisenberg and co-workers crystallized oligomers produced from peptide fragments of several disease-related amyloid proteins including Aβ and found out-of-register anti-parallel β-sheets in oligomers and fibrils [62, 63]. A recent solid state NMR study by Ishii and co-workers [64] of an Aβ(1-42) oligomer called an SPA revealed a structure with some similarities to our 150 kDa oligomer, including aggregate dimensions, predicted β-strand regions, and ^13^C linewidths (3-4 ppm) on NMR spectra. However, fpRFDR-CT NMR measurements of SPA selectively labeled at ^13^CO of A30, L34, or V39 indicated out-of-register parallel β-sheets, in contrast to the C-strand structure in our 150 kDa oligomers. These studies either didn’t have enough data to reveal all the secondary structures in the oligomers or only focused on segments of Aβ peptides. However, they did provide evidence for the existence of diverse assembly pathways for different secondary structures that lead to oligomer, an idea which had also been proposed by some simulation studies [65]. Our study is the first to demonstrate the coexistence of out-of-register parallel β-sheets and anti-parallel β-sheets within a single oligomer formed by non-modified Aβ(1-42) peptide.

The coexistence of two different β-strand alignments in 150 kDa oligomers is unexpected but also sheds more light on the full structure of the oligomers. For example, an arrangement alternative to that in Figure 8B could involve the stacking of two separate β-sheets, namely a fully anti-parallel C-strand sheet (Figure S5A, left) and an out-of-register parallel N-strand sheet (Figure 8A). Domain swapping of N- and C-strands could be further introduced to expand the possible arrangements of the two β-sheets. Finally, a possible 150 kDa oligomer structure with different alignments of β-sheets and domain swapping might suggests that the N-strand and the C-strand form β-sheets in different stages of oligomer aggregation.

## Materials and Methods

### Chemicals and Aβ(1-42) peptide synthesis

Chemical reagents used in preparation of oligomer samples were purchased from Sigma-Aldrich (St. Louis, MO). Aβ(1-42) peptides (sequence DAEFR HDSGY EVHHQ KLVFF AEDVG SNKGA IIGLM VGGVV IA) with or without ^13^C and ^15^N labels were synthesized by New England Peptide (Gardner, MA) and by the Proteomics Core at the Mayo Clinic (Rochester, MN), which both equipped Liberty Blue peptide synthesizer from CEM (Matthews, NC). The isotope-labeled compounds used in syntheses were all purchased from Cambridge Isotope Laboratories (Tewksbury, MA).

### Aβ(1-42) 150kDa oligomer preparations for solid-state NMR

Crude product of Aβ(1-42) peptide was directly treated with Size-exclusion chromatography (SEC) to isolate Aβ(1-42) monomers as previously described [22, 23]. Aliquots of SEC-purified Aβ monomer were incubated overnight with 50 mM sodium chloride and 4 mM SDS at room temperature. We got some initial small oligomers, namely 2-4mers, in the incubation [22]. The solution of 2-4mers was then dialyzed against 20 mM NaP for 48–72 h and then against 10 mM NaP for an additional 3–4 h with at least five buffer changes. SDS was removed and the concentration of salt was reduced during the dialyses. The quality of oligomer samples was tested by CD and ThT fluorescence. Finally, residual or unassembled monomers needed to be removed by filtering with an Amicon Ultra 4 centrifugal concentration/filtration device, which has a molecular mass cutoff of 50 kDa. More detailed procedure of preparing 150 kDa oligomers can be found in our previous reports [22, 23].

For solid-state NMR experiments, at least five preparations were performed for each sample to provide sufficient amounts of oligomers. The preparations for one sample were combined, flash-frozen, and immediately lyophilized. The lyophilized oligomer samples were stored at -80 °C until use. The isotope-diluted samples, e.g. Sample I, were prepared from a mixture of monomers, which was composed of isotope-labeled and unlabeled Aβ(1-42) in desired ratio.

### Solid-state NMR experiments

All the solid-state NMR experiments were performed on a Bruker narrow-bore 11.7 Tesla magnet (^1^H frequency of 500 MHz), equipped with a 3.2-mm HCN MAS probe and a 3.2-mm double-resonance MAS probe. The 2D fpRFDR [24] and 2D DARR [32, 33] spectra are 2D ^13^C-^13^C exchange experiments with different mechanisms to reintroduce dipolar coupling between ^13^C and thus providing cross-peaks. Proton decoupling with a ^1^H radiofrequency field of 100 kHz was used in fpRFDR recoupling periods and acquisitions, and two-pulse-phase modulation (TPPM) [66] was selected to be the decoupling method. In 2D DARR experiments, continuous irradiation with powers corresponding to 11 kHz nutation frequencies (same as MAS spinning rate) in ^1^H channel was applied during the exchange periods. The lengths of exchange periods were set to 50 ms or 500 ms for verifying intra-residue contacts or detecting inter-residue long-distance contacts, respectively. As for 2D fpRFDR experiment, the power of the π pulse on ^13^C channel was adjusted to be 33 kHz to match the duration (15.2 μs) of one-third of rotor period at 22 kHz MAS. The signal averaging of 2D fpRFDR and 2D DARR required 36 to 48 h to produce spectra in Figure S1, S2 and S13 to S16. For isotope-diluted samples, the signal averaging should be increased to 72 h due to less ^13^C in the sample. To determine the positions and the linewidths of crosspeaks on 2D spectra, non-linear fitting of 3D gaussian function was performed for each crosspeak.

PITHIRDS-CT experiments [28] was performed on sample A to I with MAS spinning rate of 12.5 kHz. The dipolar recoupling time was adjusted by number of blocks of pulses (k1, k2 and k3 defined by Tycko [28]), and it was fixed to be between 0 and 61.4 ms in our measurements. 100 kHz proton decoupling was conducted by continuous wave decoupling during PIRHIRDS recoupling and acquisition. PITHIRDS curves in Figure 3A and 6A were from signal averaging of about 24 hours. All the PITHIRDS data sets were corrected by subtracting background signals of natural abundance ^13^C in Aβ(1-42) molecule. We estimate the signal from natural abundance ^13^C by simply counting the number of ^13^C sites with similar chemical shifts. For backbone ^13^CO labels, there are 35 similar CO sites (excluding all glycines and the C-terminus). For alanin ^13^Cβ labels, there are 22 similar methyl sites.

### Molecular modeling

Single β-strand was generated by setting the backbone torsion angles to fixed values (Ψ=112°, ϕ=-119°). Then, the generated β-strands were translated and rotated to ensure the hydrogen bond pattern on backbones of parallel or anti-parallel β-sheet. The sidechains were kept in their initial conformations. For the idealized β-sheet models, no energy minimization or optimization was performed.

### NMR-related spin simulations

Simulated PITHIRDS-CT curves were generated using SPINEVOLUTION [67] with the use of parameters that matched the experimental conditions. Briefly, all ^13^C atoms were treated as identical spins and their positions are fixed by atom coordinates. All the initial spin vectors were in +x direction, and they evolved according to the pulse sequence of PITHIRDS. The intensities of detected signal at different time points were stored and were used to plot the simulation curves. The REPULSION powder averaging scheme was used for the simulations [68].

Simulations for singly labeled samples (Figure 3) were based on linear 8-spin system, which is a linear array of 8 ^13^C spins separated by constant distance. For doubly labeled samples, the simulated curves (Figure 7) were generated from 16-spin system, which used the coordinates of 16 ^13^CO sites from 8 strands in the idealized models of out-of-register β-sheets. For the 4-spin simulation in Figure S22B, only 4 ^13^CO sites from 2 neighboring strands are involved.

## Acknowledgements

This work was supported by the Alzheimer’s Association (grant NIRG-10-173755 to A.K.P.), the National High Magnetic Field Laboratory User Collaboration Research Grant Program, and the National Institute on Aging of the National Institutes of Health (award number R01AG045703). We gratefully acknowledge Evan K. Roberts for helping us analyzing 2D NMR spectra and proposing possible structures.

## Footnotes

1 Although DMSO was introduced to insure that initial synthetic Aβ(1-42) solutions remain disaggregated [55], it actually induces aggregate formation. Treatment of Aβ(1-42) films with DMSO and dilution into aqueous buffer [56] followed by SEC showed an increase in the amount of the aggregate peak near the void volume relative to that of Aβ(1-42) samples which had not been exposed to DMSO (T. L. Rosenberry, unpublished observations).

