## Supplementary Figures and Tables for "Out-of-register parallel β-sheets and antiparallel β-sheets coexist in 150 kDa oligomers formed by Aβ(1-42)"

Table S1. NMR chemical shifts (ppm)/linewidths (full width half maximum, ppm) for all ^13^C-labeled sites in the Aβ(1-42) 150kDa oligomer samples. The carbons in each amino are labeled as they are in the Biological Magnetic Resonance Data Bank [1]. Estimated error is ±0.1 ppm for both the chemical shift and line width unless specified otherwise. The fitting of crosspeaks is described in Materials and Methods section.

| Residue | CO | C^α^ | C^β^ | C^γ^ | Other Carbons |
| --- | --- | --- | --- | --- | --- |
| D7 | 172.7/2.8 | 51.1/2.3 | 40.0/4.8 | 176.9/3.8 |  |
| S8 | 171.6/3.3 | 56.5/4.6 | 62.3/5.2 |  |  |
| G9 | 170.8/4.7 | 43.5/3.2 |  |  |  |
| Y10 | 171.9/2.7 | 57.7/3.8 | 35.7/5.1 | 127.9/2.4 | C^δ^: 130.6/3.2,  C^ε^: 116.0/3.6,  C^ζ^: 156.3/3.1 |
| E11 | 173.8/4.5 | 53.0/3.1 | 31.7/8.1 | 34.4/3.8 | C^δ^: 180.2/3.1 |
| V12 | 174.1/3.2 | 58.8/4.0 | 32.7/2.9 | 19.3/3.3 |  |
| H13 | 171.8/1.9 | 52.5/2.3 | 31.1/2.6 | 128.8/5.3 | C^δ^: 114.7/4.4,  C^ε^: 135.9/2.8 |
| H14 |  |  |  |  |  |
| Q15 | 172.4/3.1 | 52.9/3.3 | 30.2/4.2 | 32.1/5.3 | C^δ^: 176.3/5.1 |
| K16 | 171.3/3.7 | 53.2/2.6 | 34.4/5.2 | 23.9/2.2 | C^δ^: 28.0/3.0,  C^ε^: 40.1/2.2 |
| L17 | 172.4/2.4 | 51.9/2.9 | 44.0/4.7 | 25.1/2.1 | C^δ1^: 22.4/2.7,  C^δ2^: 20.6/1.8 |
| V18 | 172.0/3.4 | 58.6/3.5 | 33.0/3.8 | 19.5/3.5 |  |
| F19 | 172.1/2.7 | 54.2/3.9 | 40.8/2.2 | 136.6/3.2 | C^δ^: 129.0/2.6 |
| F20 | 172.1/5.0 | 54.2/3.9 | 40.1/5.5 | 137.2/3.1 | C^δ^: 128.7/4.2 |
| A21 | 174.3/3.4 | 48.7/3.1 | 20.9/5.1 |  |  |
| E22 | 172.4/3.1 | 52.9/3.1 | 31.5/6.4 | 34.2/7.5 | C^δ^: 180.5/5.0 |
| D23 | 173.6/4.2 | 51.6/3.2 | 40.7/8.2 | 177.8/4.4 |  |
| V24 | 172.1/2.3 | 57.6/3.0 | 33.1/2.6 | 19.2/3.7 |  |
| G25 | 171.0/4.7 | 43.4/3.7 |  |  |  |
| S26 | 172.9/4.4 | 56.3/5.3 | 62.5/4.6 |  |  |
| N27 | 173.9/5.3 | 51.6/4.8 | 37.5/7.5 | 174.8/4.2 |  |
| K28 | 174.7/6.6 | 55.3±0.2/5.9 | 34.0/5.1 | 23.3/4.6 | C^δ^: 27.9/3.2  C^ε^: 40.7/2.6 |
| G29 | 171.0/5.0 | 43.6/3.9 |  |  |  |
| A30 | 173.1/2.0 | 48.8/2.8 | 19.9/3.8 |  |  |
| I31 | 172.1 /2.3 | 58.3/2.9 | 39.3/1.2 | Cγ1: 25.3/2.5,  Cγ2: 15.0/3.6, | C^δ^: 12.0/2.1 |
| I32 | 170.6/2.1 | 57.5/2.9 | 39.7/3.1 | Cγ1: 25.6/2.8, Cγ2: 15.9/3.7 | C^δ^: 12.3/2.4 |
| G33 | 169.3/2.1 | 43.1/2.8 |  |  |  |
| L34 | 171.1/3.7 | 51.6/2.3 | 44.4/2.8 | 24.5/2.5 | C^δ^: 21.6±0.4/1.8±0.3 |
| M35 | 171.8/2.1 | 52.7/2.4 | 35.4/3.5 | 30.4/1.6 | C^ε^: 15.6/2.3 |
| V36 | 172.4/3.0 | 57.6/3.1 | 33.6/2.9 | 19.3/2.9 |  |
| G37 | 169.3/1.9 | 43.8/2.7 |  |  |  |
| G38 | 168.5/1.8 | 44.0/2.2 |  |  |  |
| V39 | 171.9/3.1 | 57.8/2.7 | 34.0/2.5 | 19.5/2.7 |  |
| V40 | 172.5/2.5 | 58.8/3.2 | 33.1/3.0 | 19.4/2.9 |  |
| I41 | 171.3/3.3 | 58.6/2.9 | 37.6/4.5 | C^γ1^: 25.5/2.2,  C^γ2^: 15.7/3.0, | C^δ^: 12.2/2.5 |
| A42 |  |  |  |  |  |

Table S2. The TALOS-N prediction of torsion angles based on ^13^C chemical shifts [2]. The blank rows indicate there is no reliable prediction for some residues. The ϕ and ψ are the predicted backbone torsion angles (defined in Reference [3]), and the β-strand like torsion angles are highlighted in green. The Dϕ and Dψ are the estimated standard deviations of the prediction errors. The S2 is the Wishart RCI chemical shift order parameter [4] that indicate the flexibility of the conformation. The secondary structure prediction is listed in one letter code: H: Helix, E: Strand, L: Coil.

| **Res Num** | **Res Type** | **ϕ (deg)** | **ψ (deg)** | **Dϕ (deg)** | **Dψ (deg)** | **S2** | **Secondary Structure Prediction** |
| --- | --- | --- | --- | --- | --- | --- | --- |
| 7 | D | -95.446 | 37.157 | 12.043 | 44.175 | 0.653 | L |
| 8 | S | -64.102 | 138.893 | 9.916 | 9.462 | 0.673 | L |
| 9 | G | 114.514 | 13.817 | 64.417 | 49.236 | 0.719 | L |
| 10 | Y | -121.556 | 151.202 | 36.343 | 18.782 | 0.789 | L |
| 11 | E | -126.595 | 147.437 | 13.754 | 11.349 | 0.835 | E |
| 12 | V | -108.119 | 135.498 | 14.987 | 12.909 | 0.825 | E |
| 13 | H | -126.207 | 146.346 | 17.363 | 13.571 | 0.813 | E |
| 14 | H | -75.993 | 138.076 | 11.516 | 9.017 | 0.819 | E |
| 15 | Q | -130.142 | 149.664 | 18.961 | 11.456 | 0.879 | L |
| 16 | K | -126.600 | 145.660 | 13.151 | 12.544 | 0.892 | E |
| 17 | L | -122.132 | 131.743 | 13.309 | 7.272 | 0.911 | E |
| 18 | V | -112.984 | 129.792 | 10.359 | 8.528 | 0.910 | E |
| 19 | F | -112.285 | 128.259 | 9.291 | 10.324 | 0.905 | E |
| 20 | F | -119.588 | 136.351 | 12.716 | 11.738 | 0.898 | E |
| 21 | A | -131.677 | 147.280 | 13.672 | 11.378 | 0.887 | E |
| 22 | E | -134.064 | 144.045 | 12.725 | 10.330 | 0.872 | E |
| 23 | D | -101.792 | 127.653 | 17.477 | 12.416 | 0.847 | E |
| 24 | V | -131.628 | 148.591 | 10.246 | 6.609 | 0.820 | E |
| 25 | G | -114.867 | 160.997 | 22.531 | 15.836 | 0.680 | L |
| 26 | S | -88.422 | 145.704 | 21.648 | 14.571 | 0.535 | L |
| 27 | N | -81.735 | 151.486 | 25.622 | 21.674 | 0.446 | L |
| 28 | K | -109.382 | 140.280 | 36.356 | 18.478 | 0.507 | L |
| 29 | G | -125.252 | 168.078 | 50.797 | 25.849 | 0.652 | L |
| 30 | V | -127.682 | 144.298 | 12.679 | 10.123 | 0.857 | E |
| 31 | I | -112.933 | 129.747 | 9.892 | 5.923 | 0.902 | E |
| 32 | I | -128.090 | 135.877 | 10.474 | 12.194 | 0.920 | E |
| 33 | G | -127.332 | 146.370 | 16.620 | 17.095 | 0.918 | E |
| 34 | L | -134.166 | 143.788 | 12.689 | 12.753 | 0.924 | E |
| 35 | M | -129.677 | 134.178 | 8.152 | 8.996 | 0.914 | E |
| 36 | V | -129.460 | 140.214 | 10.165 | 11.794 | 0.896 | E |
| 37 | G |  |  |  |  |  |  |
| 38 | G |  |  |  |  |  |  |
| 39 | V | -129.079 | 140.454 | 15.728 | 13.097 | 0.887 | E |
| 40 | V | -111.366 | 129.300 | 10.314 | 7.409 | 0.882 | E |
| 41 | I | -105.484 | 127.524 | 11.327 | 8.968 | 0.876 | E |
| 42 | A |  |  |  |  |  |  |

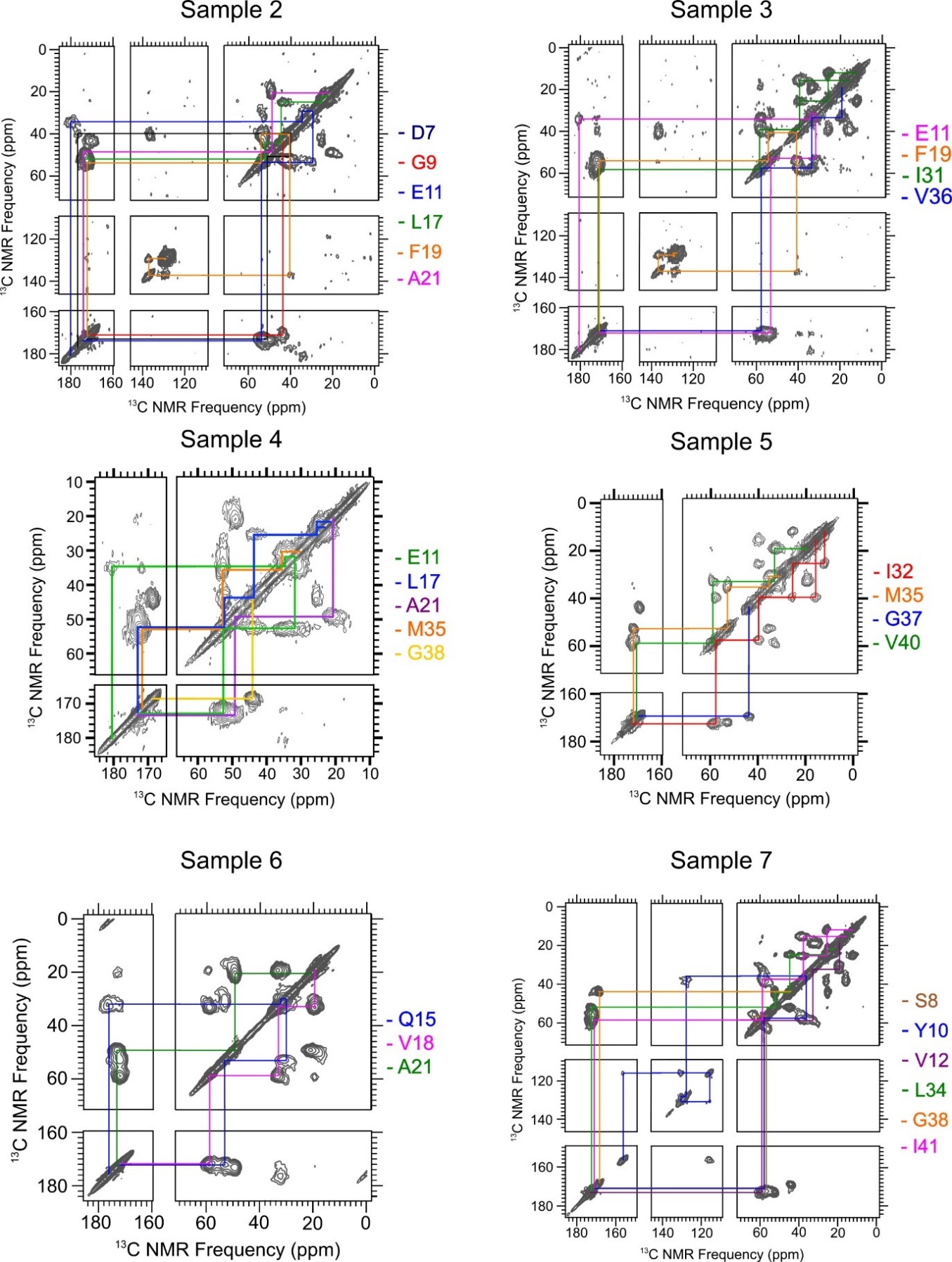

Figure S1: The fpRFDR spectra of the isotope-labeled Aβ_1-42_ 150kDa oligomers from Samples 2 to 7 in Table 1.

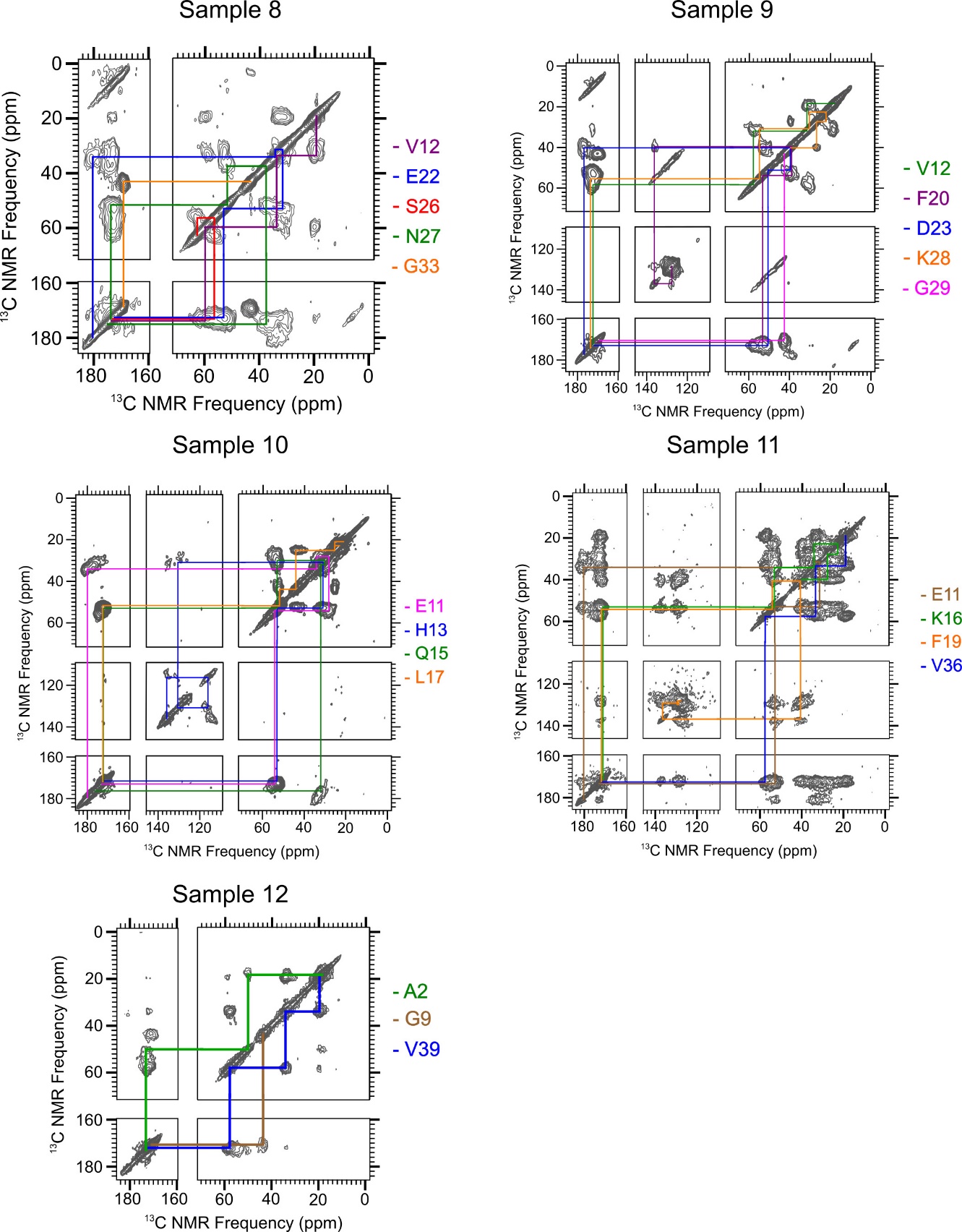

Figure S2: The fpRFDR spectra of the isotope-labeled Aβ_1-42_ 150kDa oligomers from Samples 8 to 12 in Table 1.

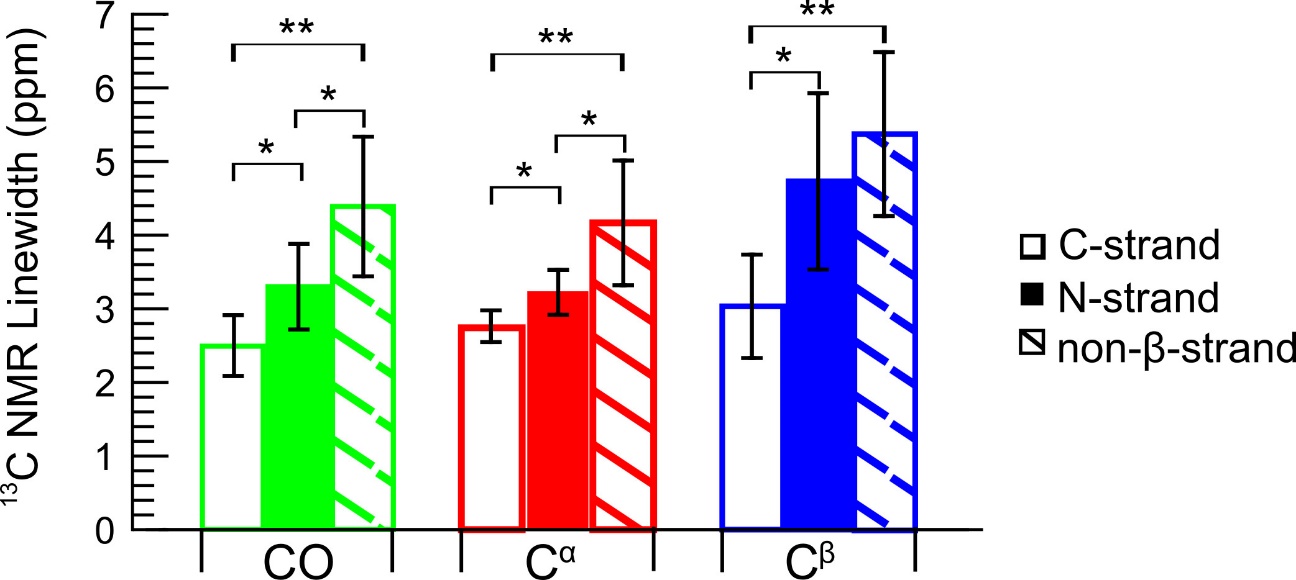

Figure S3: Measured average ^13^C NMR linewidths (full width half maximum), with error bars indicating 95% confidence intervals for CO, C^α^, and C^β^ NMR peaks measured for residues within the N-strand, C-strand, and non-β-strand regions predicted by the TALOS-N software (see Figure 2A). The Student’s t-test indicated that CO sites exhibited a significantly different average line width (P<0.05) for C-strand residues, N-strand residues, and residues within non-β-strand regions. Comparison of the C^α^ linewidths using the t-test also revealed significant differences for N-strand, C-strand, and non-β-strand regions. Comparison of C^β^ line widths, revealed a significant difference between the average C^β^ linewidth for the N-strand the C-strand, but not between the N-strand and the non-β-strand regions. (*: P<0.05, **: P<0.01)

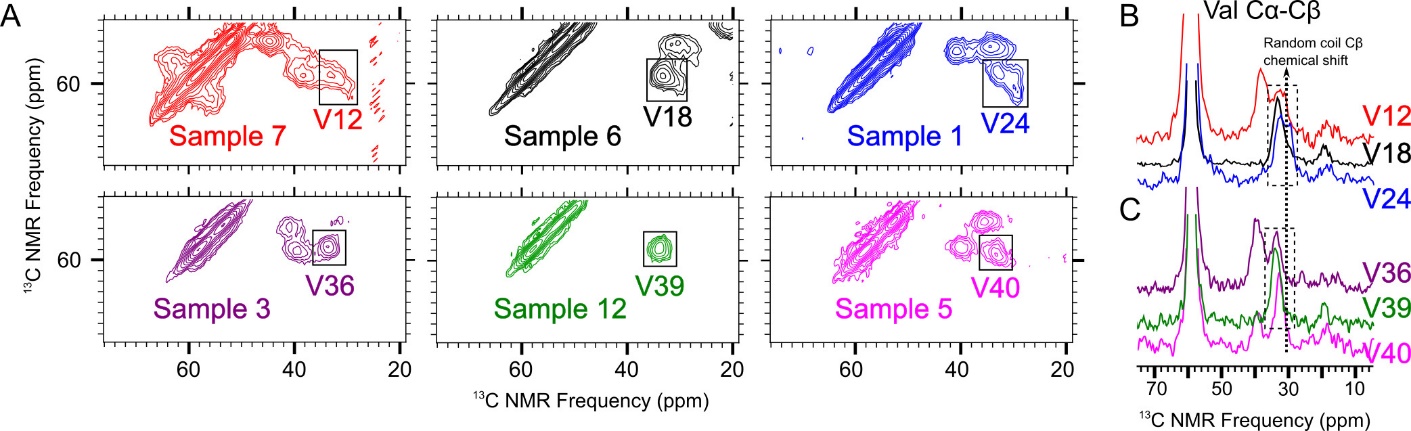

Figure S4: A) Selected regions of ^13^C-^13^C 2D-fpRFDR spectra from Samples 7, 6, 1, 3, 12, and 5, respectively, showing C^α^-C^β^ cross-peaks for all valines in the Aβ_1-42_ peptide in 150kDa oligomers. B) Slices taken at the frequency of peak intensity of the valine C^α^-C^β^ cross-peak for the spectra containing signals from V12, V18, and V24, respectively. The dashed box shows that the shape of each C^β^ signal does not correspond to a single Gaussian peak. C) Horizontal slices taken at the frequency of peak intensity of the valine C^α^-C^β^ cross-peak for the spectra containing signals from V36, V39, and V40, respectively. The dashed box shows that the shape of each C^β^ peak does correspond to single Gaussian peaks. The vertical dashed line on 31.2 ppm labels the random-coil chemical shift of Cβ in valine.

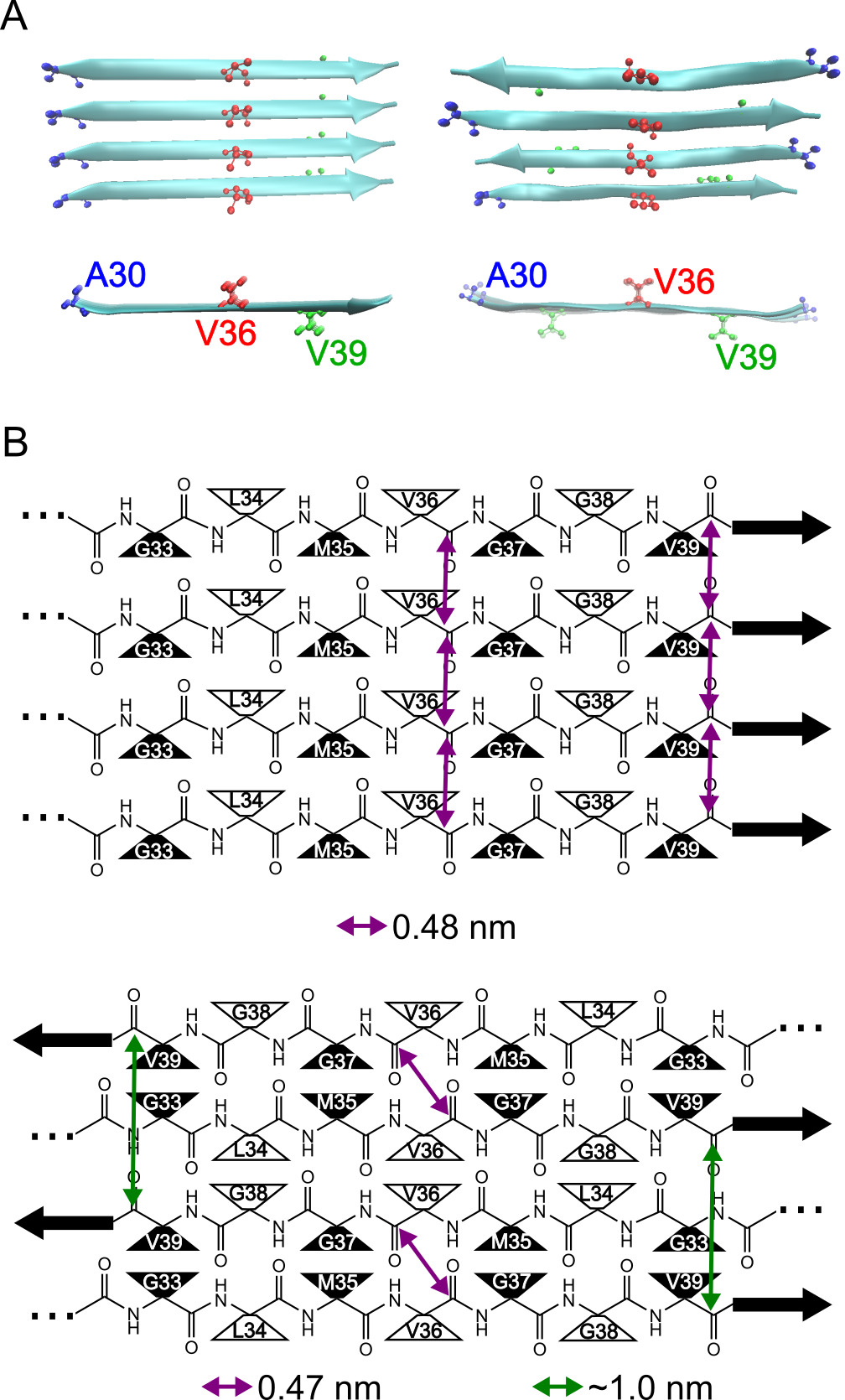

Figure S5: A) Representations of all-atom models for an in-register parallel β-sheet formed by the C-strand (left) and an antiparallel C-strand β-sheet centered at residue V36 (right). Molecular backbones are represented as blue ribbons and atoms are drawn for residues A30, V36, and V39 to indicate relative positions of ^13^C-labeled sites for the data in Figure 3. B) Schematic representations for the same β-sheet models as in Panel A (parallel β-sheet above, antiparallel β-sheet below) drawn to highlight the relative positions of backbone atoms. Sidechains are depicted as black or white triangles to illustrate the directions of residue sidechains (black: below the β-sheet plane, white: above the β-sheet plane).

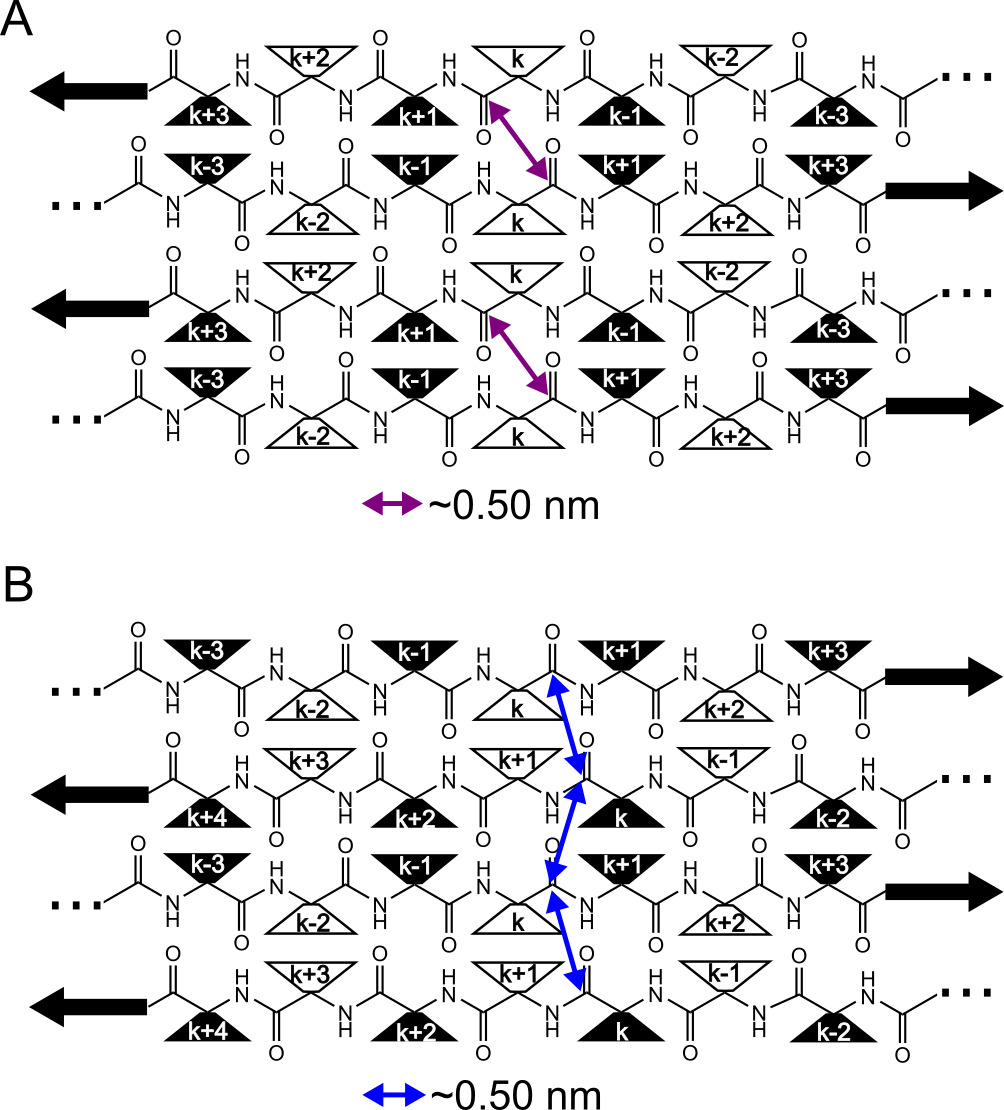

Figure S6: Schematics illustrating two possible types for anti-parallel β-sheets. Panel A shows a Type-I antiparallel β-sheet, in which sidechains of a specific residue (e.g., the k^th^ residue) are all oriented on the same face above or below the plant defined by the β-strand backbones. The sidechains are represented by black or white rectangles, indicating that the sidechain is below or above the β-sheet plane, respectively. Panel B shows a Type-II antiparallel β-sheet, in which sidechains of a specific residue (e.g., the k^th^ residue) alternate in orientation above or below the β-sheet plane between adjacent β-strands. For both Type-I and Type-II antiparallel β-sheets, there is only one central residue (the k^th^ residue in each diagram), whose carbonyl C atoms are separated by an inter-molecular distance of about 0.5 nm.

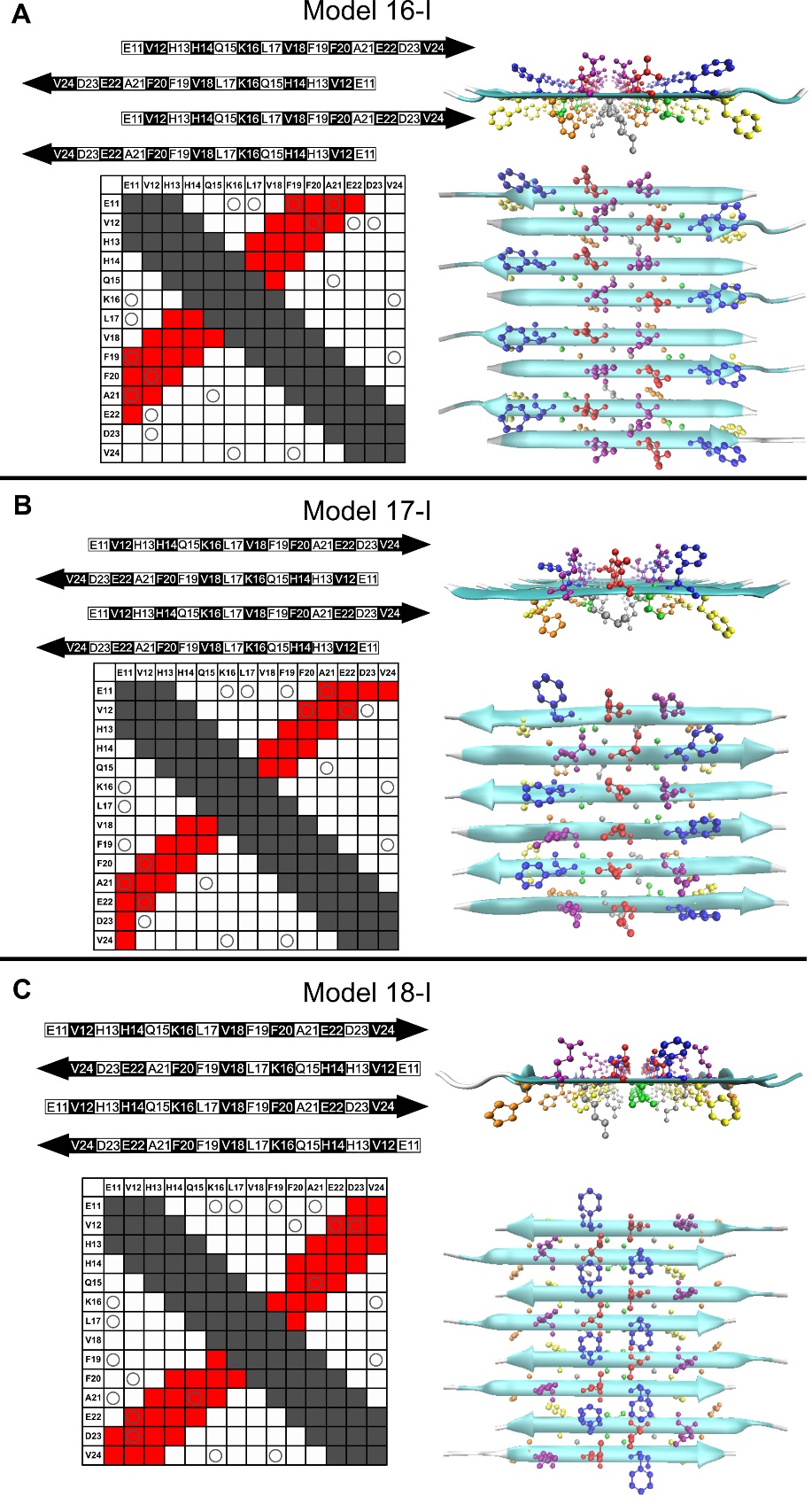

Figure S7: Possible alignments of N-strand in type-I anti-parallel β-sheet (defined in Figure S6). For each alignment, the top and side view of an idealized all atom model are listed. The color code for residues are kept identical: Orange H14, Purple Q15, Grey K16, Red L17, Green V18, Blue F19, Yellow F20. The corresponding contact charts use the same denotation as the Figure 4B.

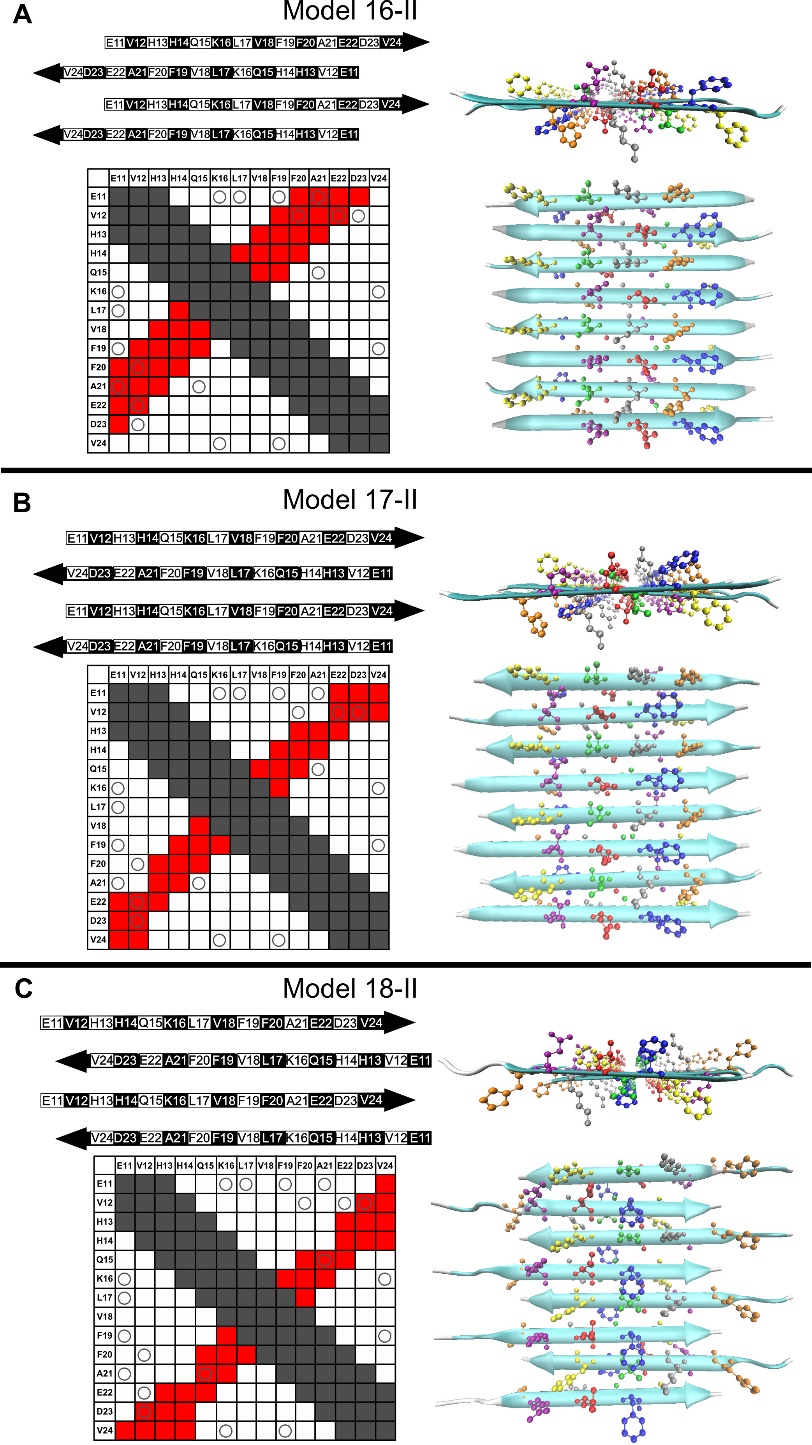

Figure S8: Possible alignments of N-strand in type-II anti-parallel β-sheet (defined in Figure S6). For each alignment, the top and side view of an idealized all atom model are listed. The color code for residues are kept identical: Orange H14, Purple Q15, Grey K16, Red L17, Green V18, Blue F19, Yellow F20. The corresponding contact charts use the same denotation as the Figure 4B.

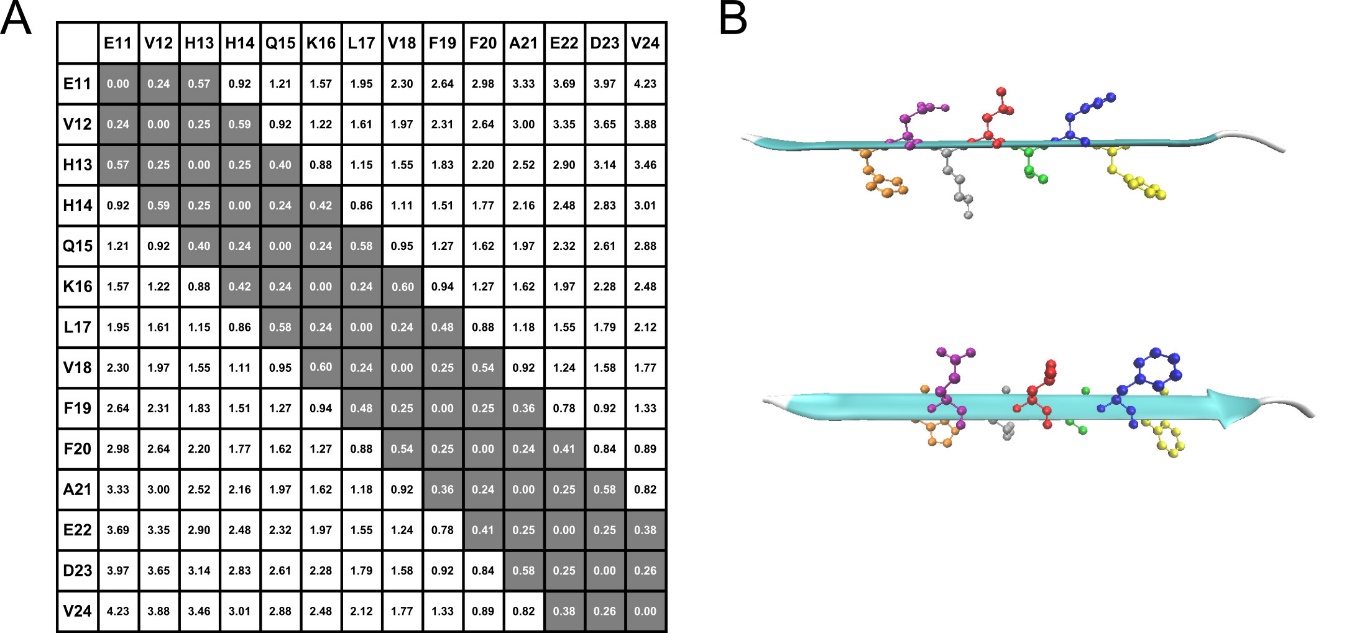

Figure S9: Residue distances between each residue within one N-strand (assuming it is β-strand conformation). A) residue distances between each residue. B) the top and side view of the all-atom model of single N-strand.

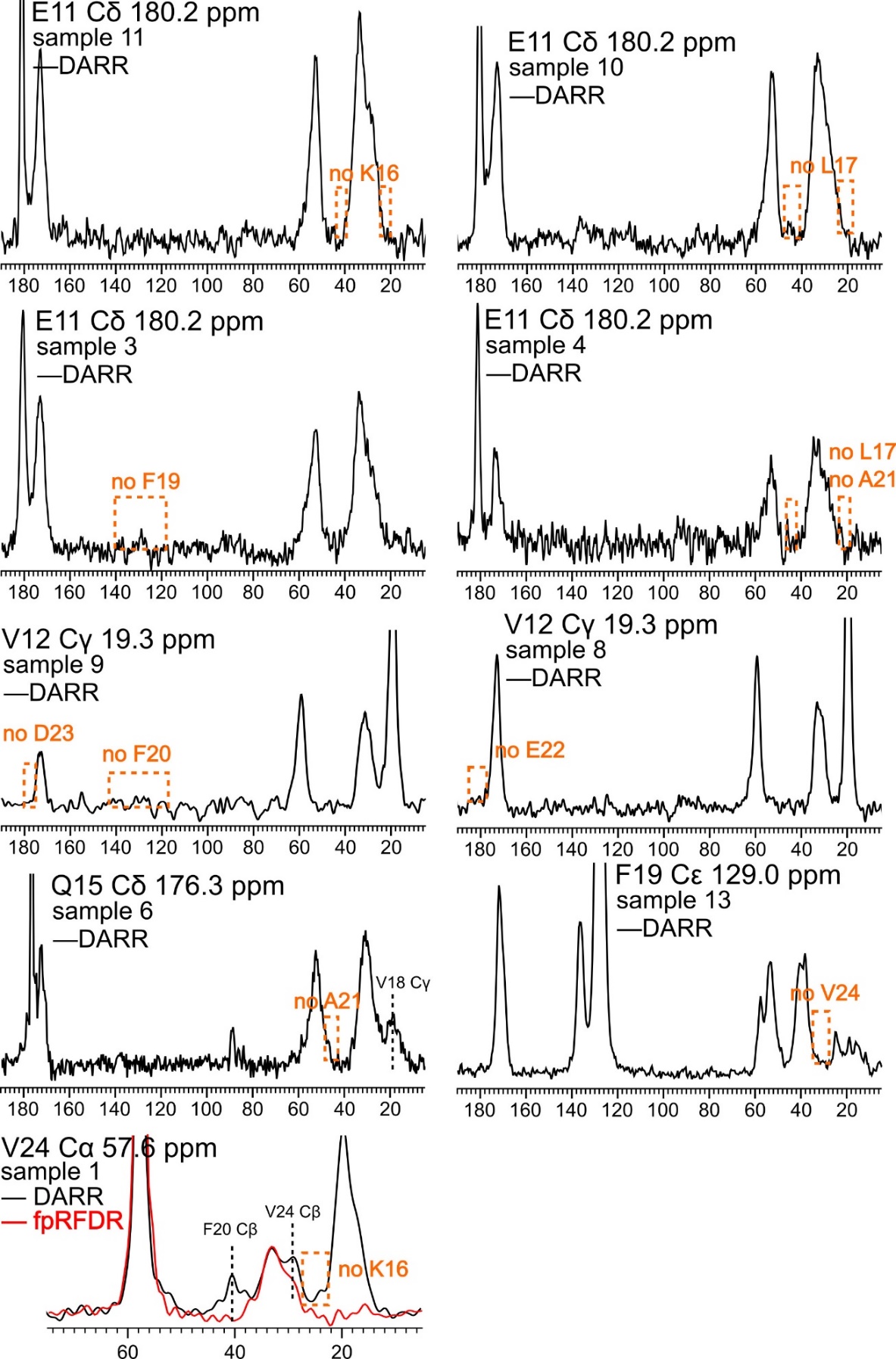

Figure S10: DARR slice with negative results from all isotope-labeled Aβ_1-42_ 150kDa oligomer samples. All the negative results in Figure 4B can be verified.

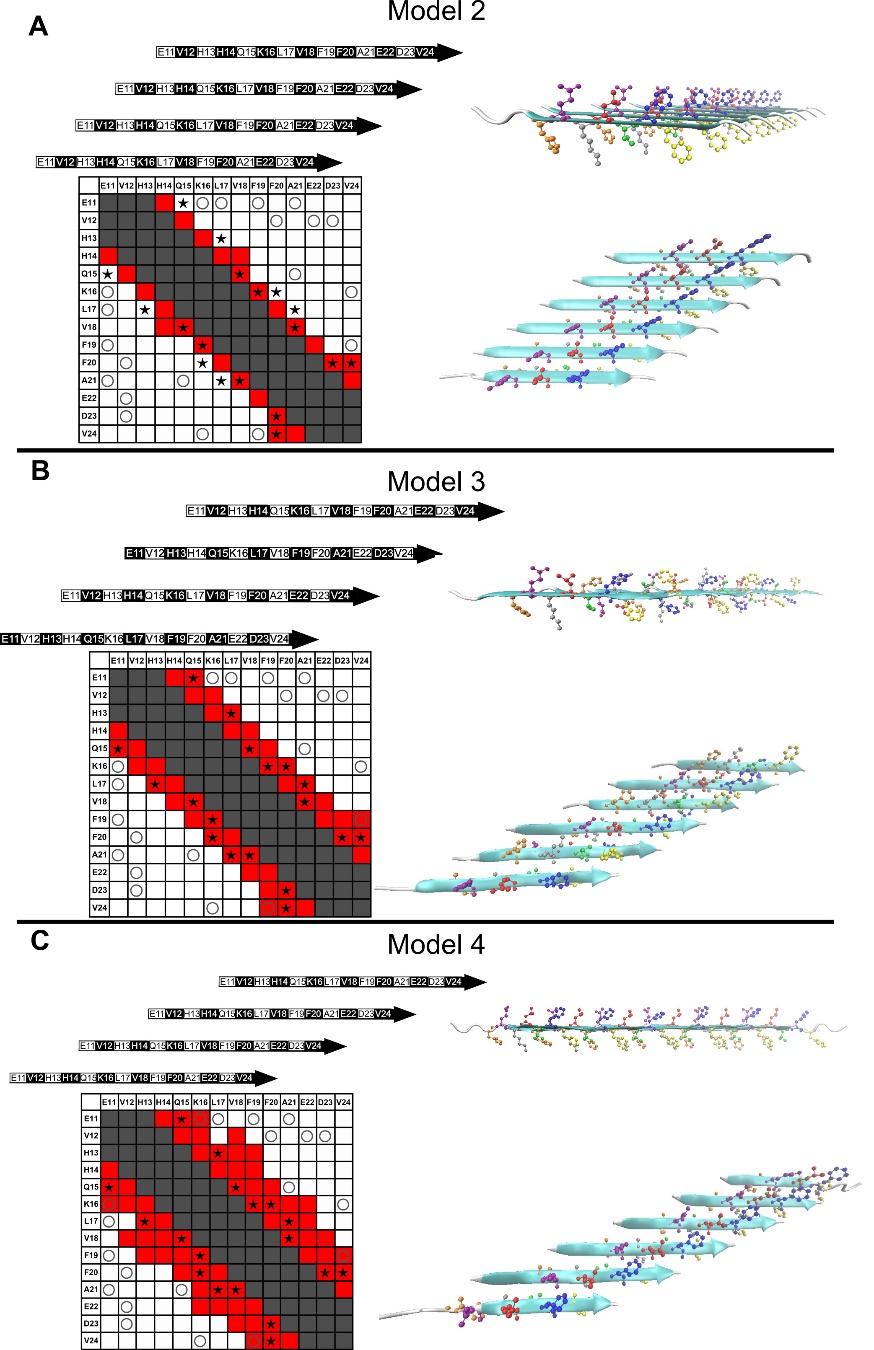

Figure S11: Possible alignments of N-strand in parallel β-sheet. For each alignment, the top and side view of an idealized all atom model are listed. The color code for residues are kept identical: Orange H14, Purple Q15, Grey K16, Red L17, Green V18, Blue F19, Yellow F20. The corresponding contact charts use the same denotation as Figure 5B.

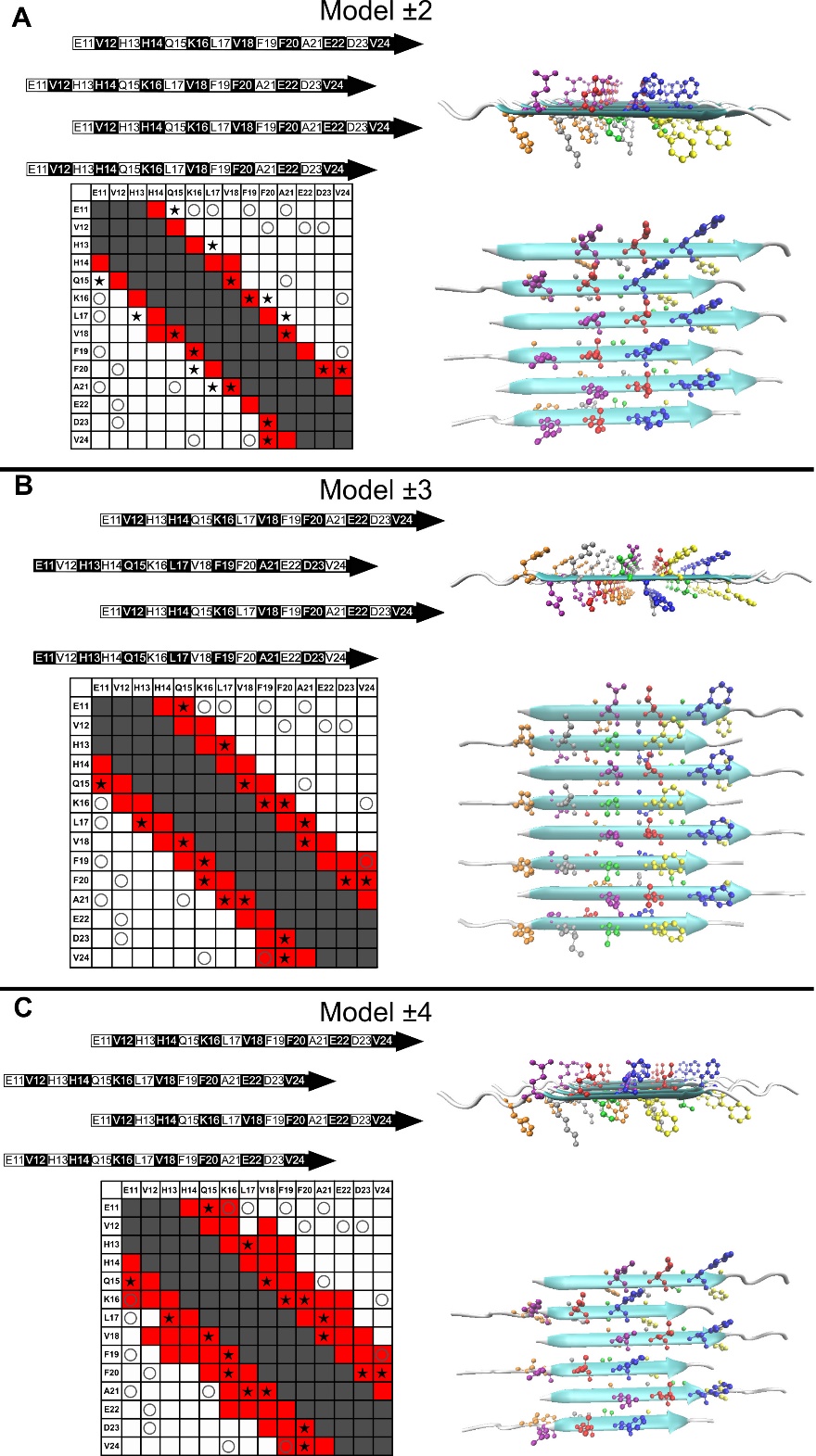

Figure S12: Possible alignments of N-strand in parallel β-sheet. For each alignment, the top and side view of an idealized all atom model are listed. The color code for residues are kept identical: Orange H14, Purple Q15, Grey K16, Red L17, Green V18, Blue F19, Yellow F20. The corresponding contact charts use the same denotation as Figure 5B.

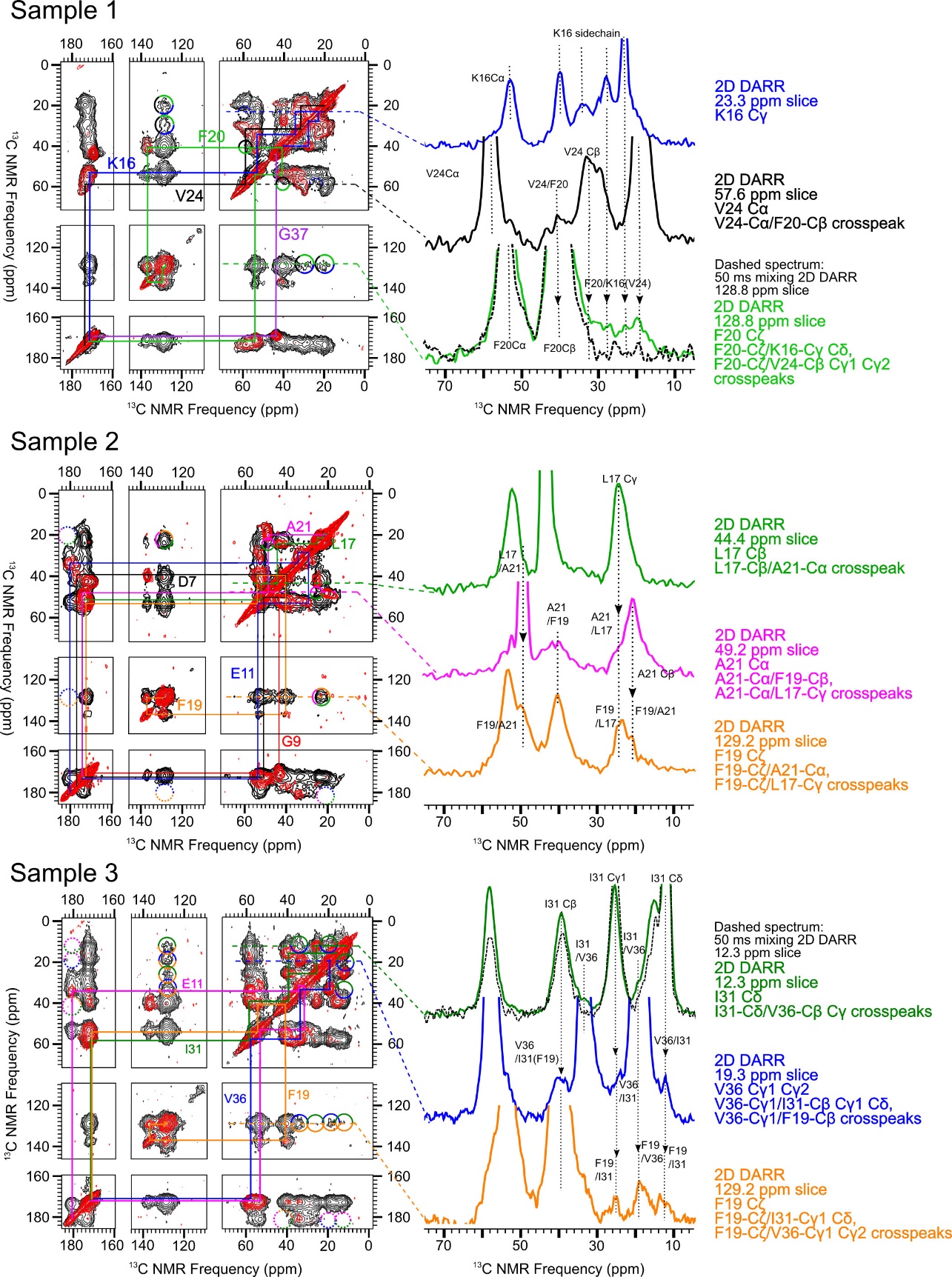

Figure S13: DARR spectra from all the isotope-labeled samples (sample 1-3). Black contours: 500ms mixing time 2D DARR spectra. Red contours: 2D fpRFDR or short mixing time 2D DARR spectra to show the intra-residue ^13^C-^13^C cross-peaks. The slices of some inter-residue cross-peaks in 500 ms mixing time 2D DARR are listed, and some slices are overlaid with the intra-residues slice (from 2D fpRFDR or short mixing time 2D DARR) to verify the ambiguous cross-peaks.

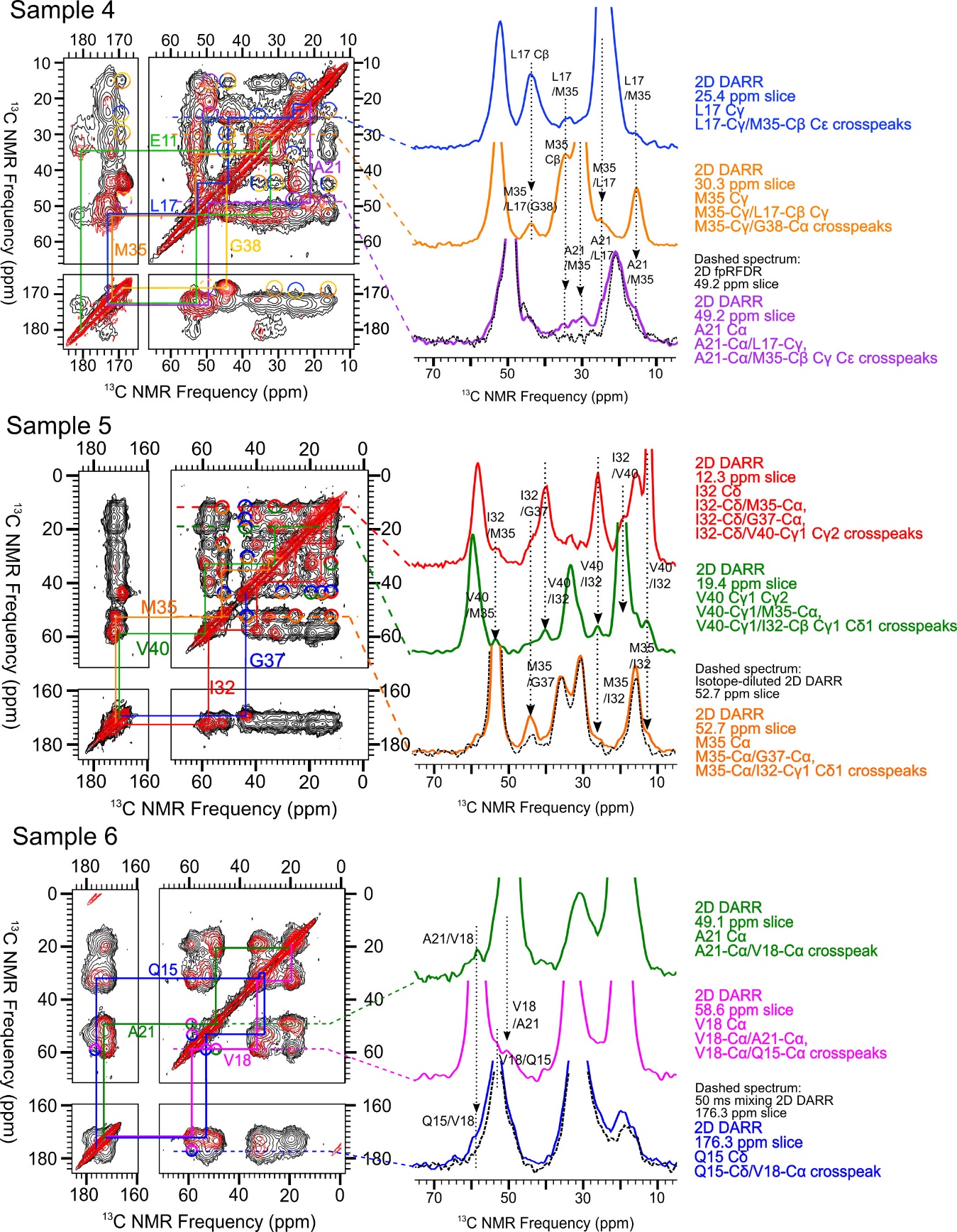

Figure S14: DARR spectra from all the isotope-labeled samples (sample 4-6). Black contours: 500ms mixing time 2D DARR spectra. Red contours: 2D fpRFDR or short mixing time 2D DARR spectra to show the intra-residue ^13^C-^13^C cross-peaks. The slices of some inter-residue cross-peaks in 500 ms mixing time 2D DARR are listed, and some slices are overlaid with the intra-residues slice (from 2D fpRFDR or short mixing time 2D DARR) to verify the ambiguous cross-peaks.

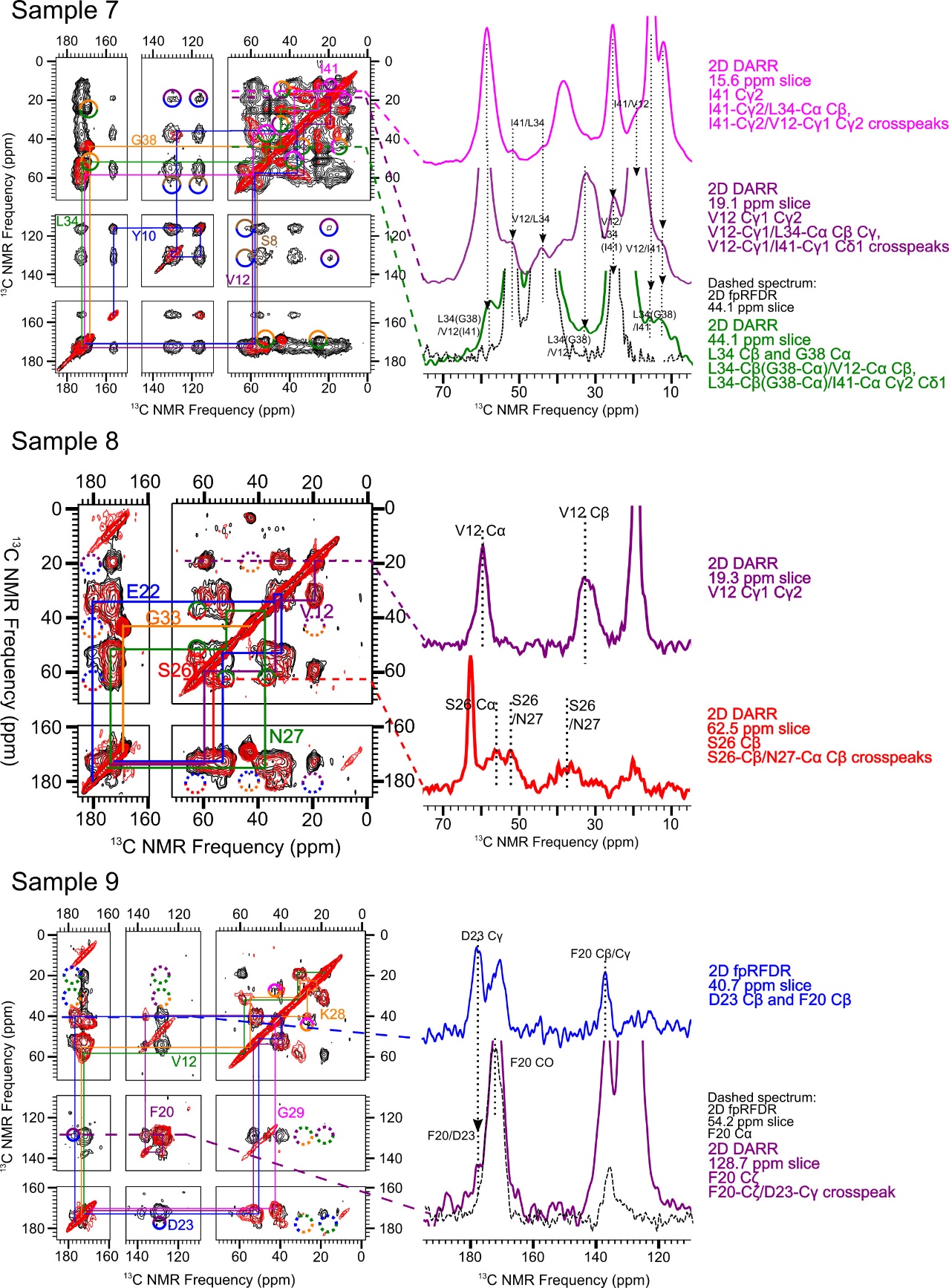

Figure S15: DARR spectra from all the isotope-labeled samples (sample 7-9). Black contours: 500ms mixing time 2D DARR spectra. Red contours: 2D fpRFDR or short mixing time 2D DARR spectra to show the intra-residue ^13^C-^13^C cross-peaks. The slices of some inter-residue cross-peaks in 500 ms mixing time 2D DARR are listed, and some slices are overlaid with the intra-residues slice (from 2D fpRFDR or short mixing time 2D DARR) to verify the ambiguous cross-peaks.

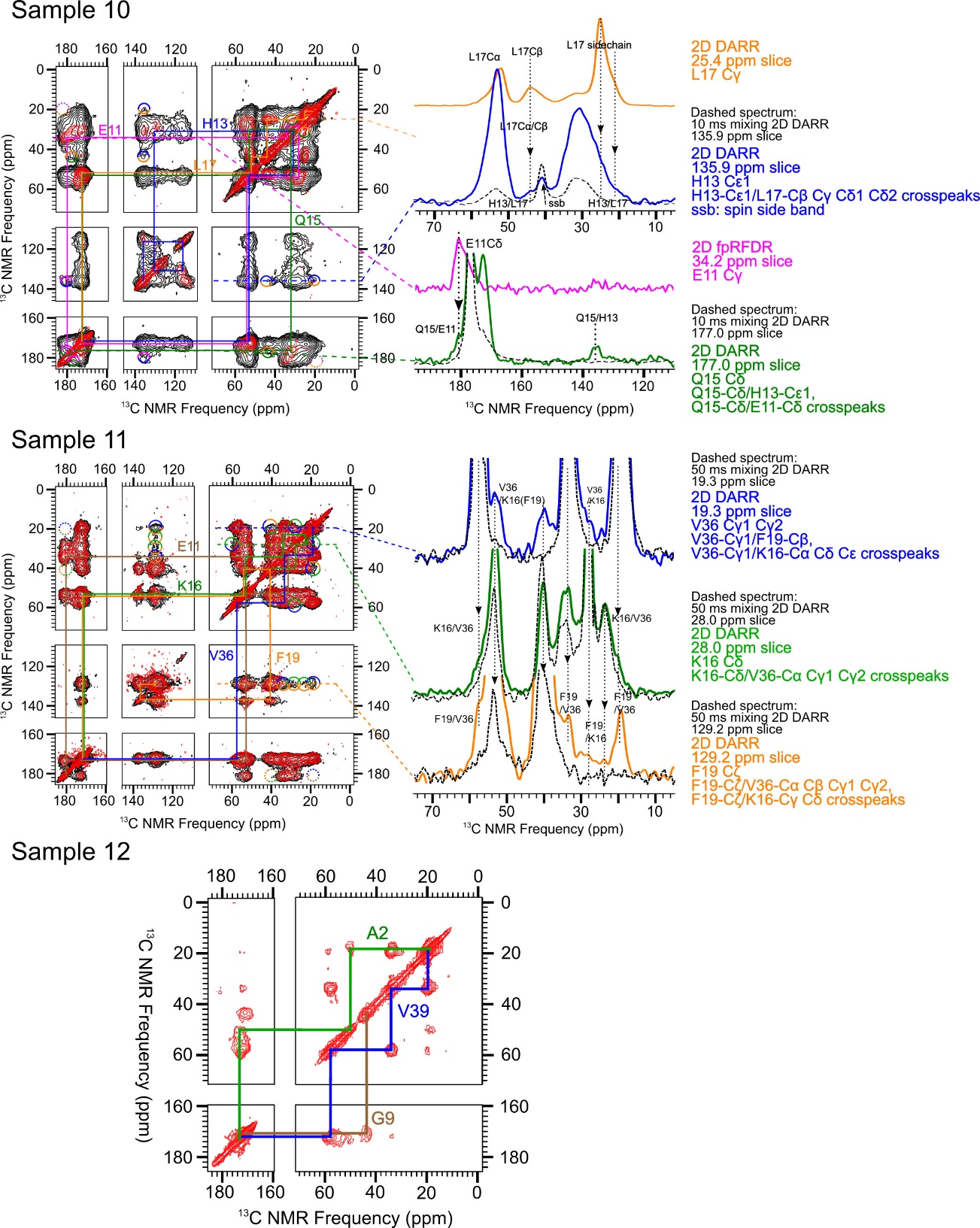

Figure S16: DARR spectra from all the isotope-labeled samples (sample 10-12). Black contours: 500ms mixing time 2D DARR spectra. Red contours: 2D fpRFDR or short mixing time 2D DARR spectra to show the intra-residue ^13^C-^13^C cross-peaks. The slices of some inter-residue cross-peaks in 500 ms mixing time 2D DARR are listed, and some slices are overlaid with the intra-residues slice (from 2D fpRFDR or short mixing time 2D DARR) to verify the ambiguous cross-peaks. Due to the low intensity of signal from A2, E3, and F4 in sample 12, the 2D DARR spectra were not collected.

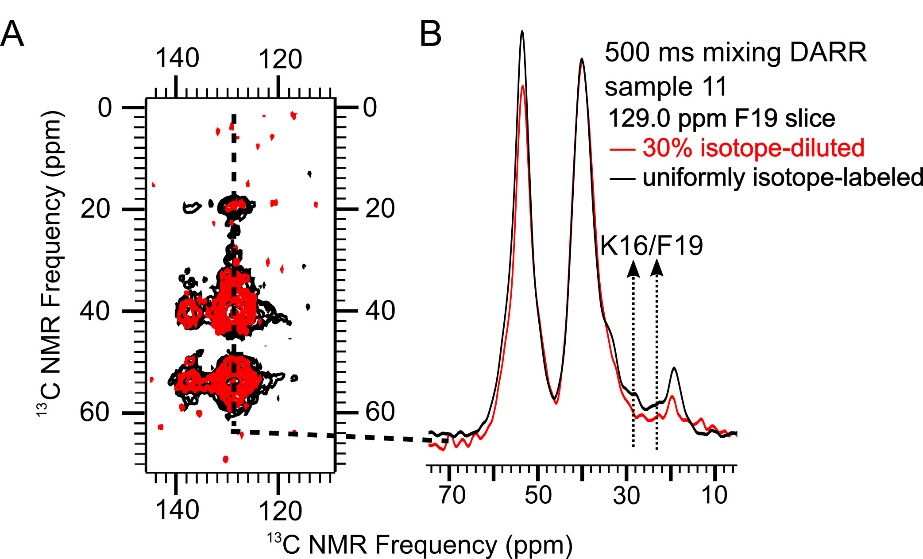

Figure S17: Isotope-dilution experiments showing that the K16/F19 contact in 150 kDa Aβ_1-42_ oligomers is inter-molecular. A) the overlaid 500ms mixing time 2D DARR spectra from uniformly isotope-labeled sample 11 (black) and the corresponding 30% isotope-diluted sample (red). B) The slices from the isotope-diluted and non-diluted 2D DARR spectra. The dilution effect of the cross-peaks between K16/F19 is labeled.

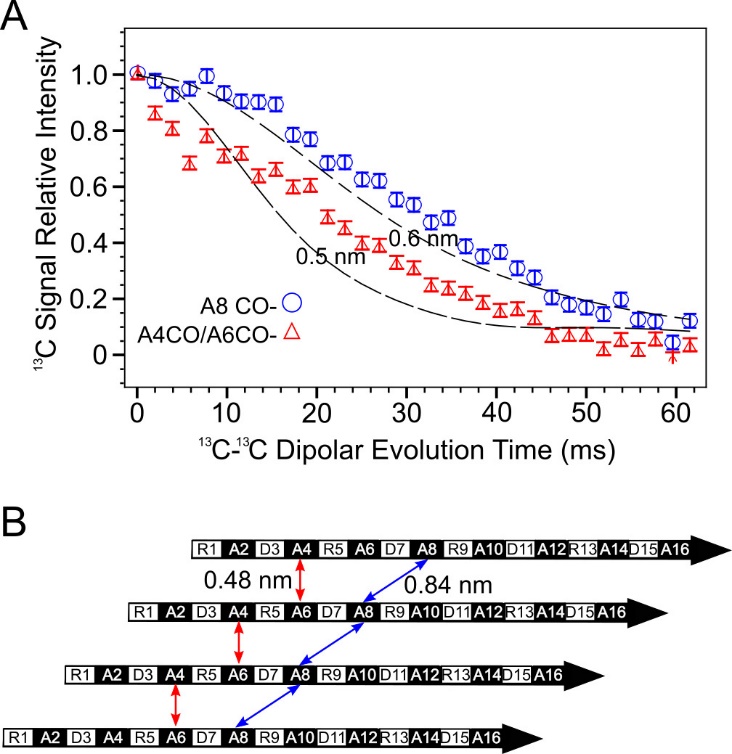

Figure S18. A) PITHIRDS-CT data for RAD16-I nanofibers, with ^13^C-labels either at the A8 CO site or the A4 CO site and the A6 CO site [5]. ^13^C-Labeling at the single A8 CO site on the β-strand backbone yielded a weak PITHIRDS-CT decay that is not consistent with in-register parallel β-sheets but is reasonable according to the short inter-sheet distance [5]. With the sample labeled at A4 and A6, we were able to detect a PITHIRDS-CT decay that is consistent with 0.48 nm nearest-neighbor ^13^C-^13^C distances. This inter-molecular distance is less than the intramolecular distance of 0.65 nm between the labeled sites. These results are consistent with a parallel β-sheet with a registry shift of 2. The dashed lines were the same simulated curves as in Figure 3. B) A schematic of a parallel β-sheet within a RADA16-I nanofiber with a registry shift of 2. Such a registry shift would place every A4 CO cite at a distance of 0.48 nm from a A6 CO cite on an adjacent strand and every A6 CO cite at a distance of 0.48 nm from a A4 CO cite on an adjacent strand. This inter-molecular distance is less than the intramolecular distance of 0.65 nm between the labeled sites. The double-headed arrows indicate distances between ^13^C-labeled CO sites.

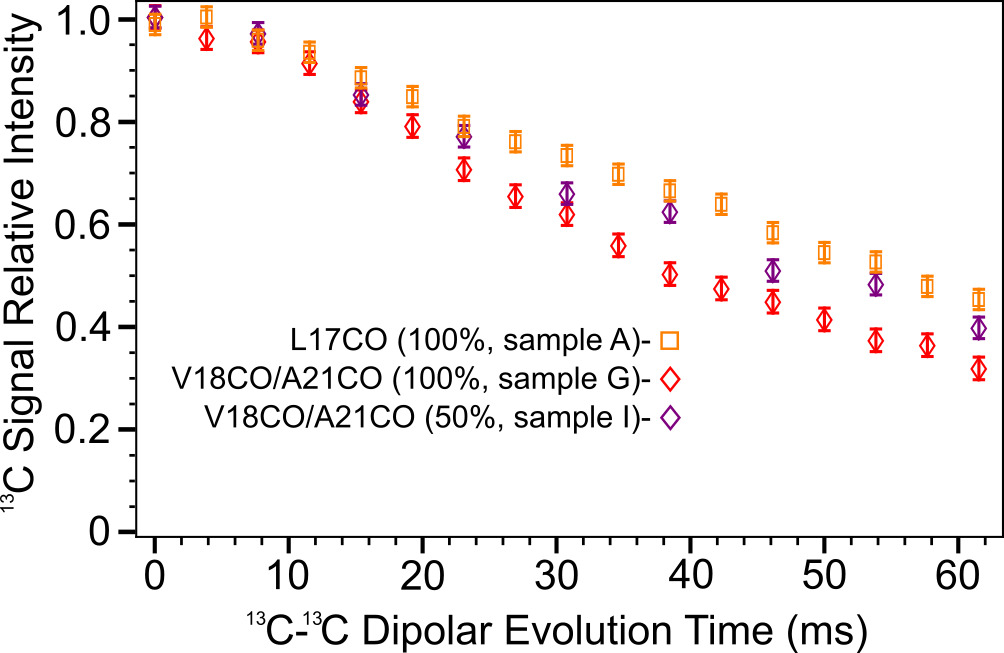

Figure S19: Isotope-dilution effect in PITHIRDS-CT data of the doubly labeled V18 CO and A21 CO sample (Sample G and I). The data from Sample A (^13^C-labeled L17CO) was also plotted here to provide the reference for the fully uncoupled ^13^C-spin system (as in Figure 3A). The PITHIRDS-CT curve of the isotope-diluted Sample I moves towards the one of singly labeled Sample A, which indicates the intermolecular coupling is dominant in doubly ^13^CO labeled samples.

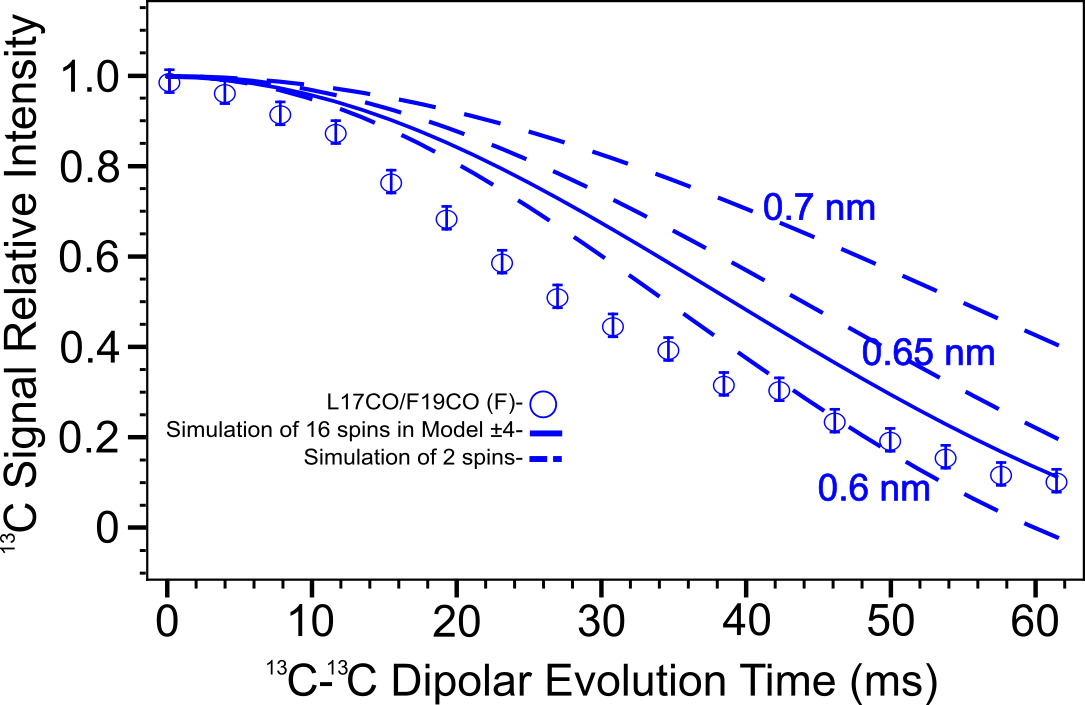

Figure S20: The PITHIRDS-CT simulated curve (solid line) of sample F in Model ±4 is similar with 2 spin simulations (dashed lines). The experimental data decay (circles) are distinct from the shape of simulated curves of two spin systems.

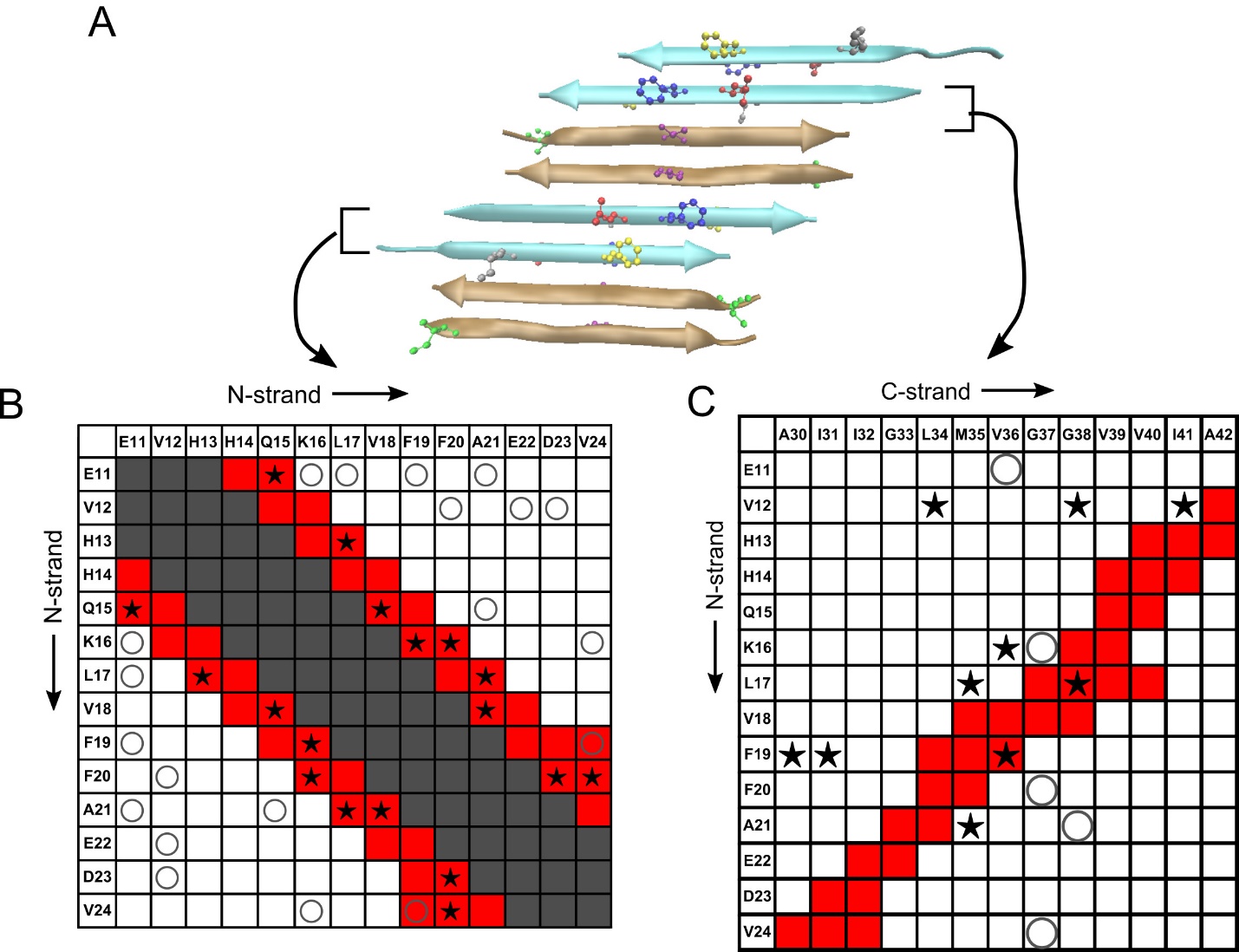

Figure S21: A) The all-atom structural model of a mixed β-sheet formed by both N-strands and C-strands. The color code of strands and residues is the same as Figure 9B. B) The contact chart of pairs of residues in neighboring N-strands. The registry shift between neighboring N-strand is 3, and the contact chart here is the same as Figure 5B. C) The contact chart that presents the observed and the predicted interactions between the neighboring N-strand and C-strand. 500 ms 2D-DARR spectra from the labeled 150 kDa oligomer samples in Table 1 were examined for crosspeaks, and inter-residue contacts were detected for 10 pairs and absent for 5 other pairs. One particular registry between N- and C-strands was analyzed, namely the, one with the observed close contact between F19 in the N-strand and V36 in the C-strand, and the contact chart shown was generated. Some observed cross-peaks between residues certainly do not match the proposed structural model (e.g. F19/I31).

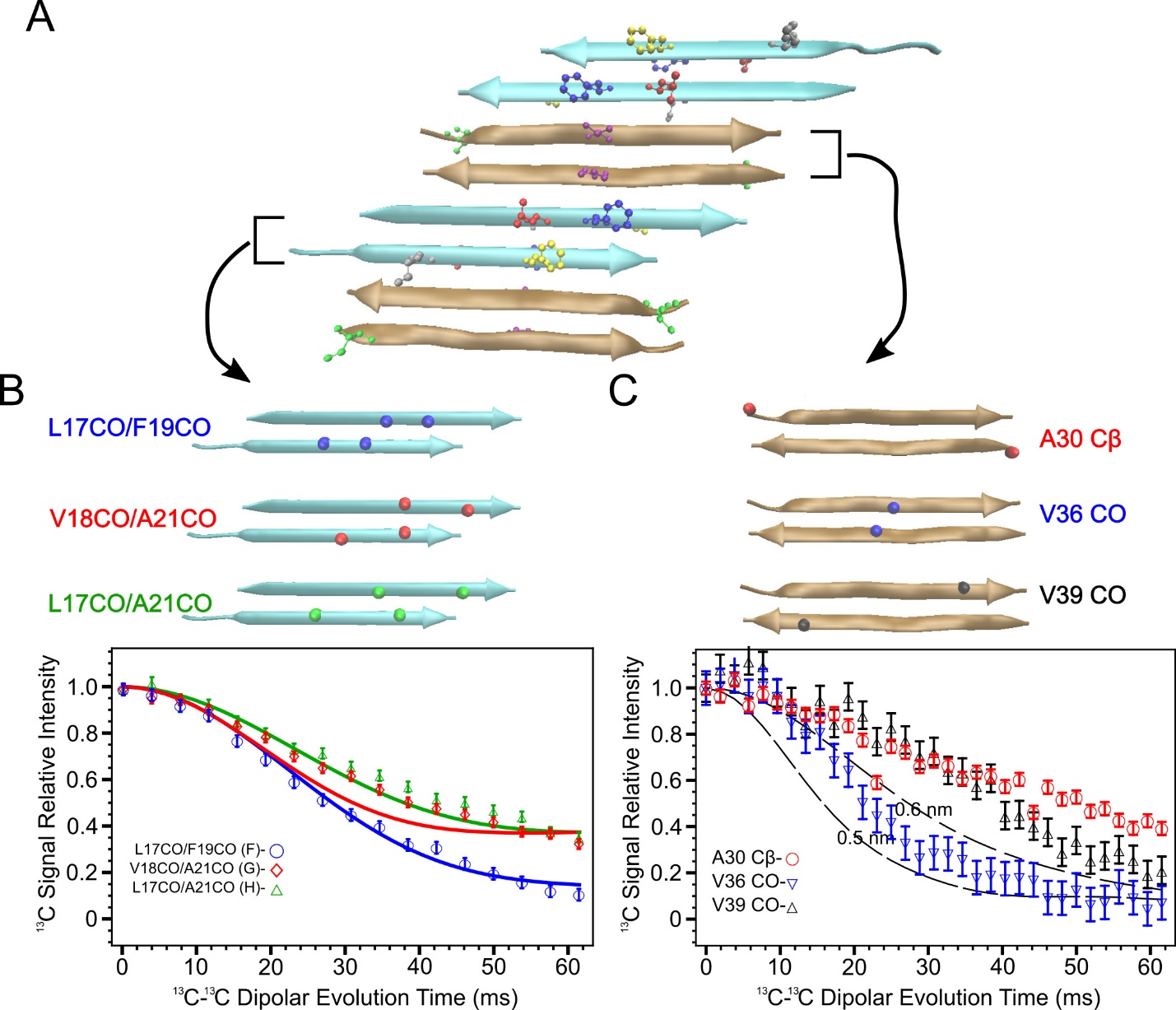

Figure S22: A) The all-atom structural model of a mixed β-sheet formed by both N-strands and C-strands. The color code of strands and residues is the same as Figure 9B. B) The PITHIRDS-CT data and the simulated curves of doubly carbonyl labeled samples according to the atom coordinates of neighboring N-strands involving only 4 ^13^C spins (Blue: sample F, Red: sample G, Green: sample H). C) The PITHIRDS-CT data of singly ^13^C-labels in C-strands (Red: A30 Cβ, Blue: V36 CO, Black: V39 CO). The same dataset is also present in Figure 3C.

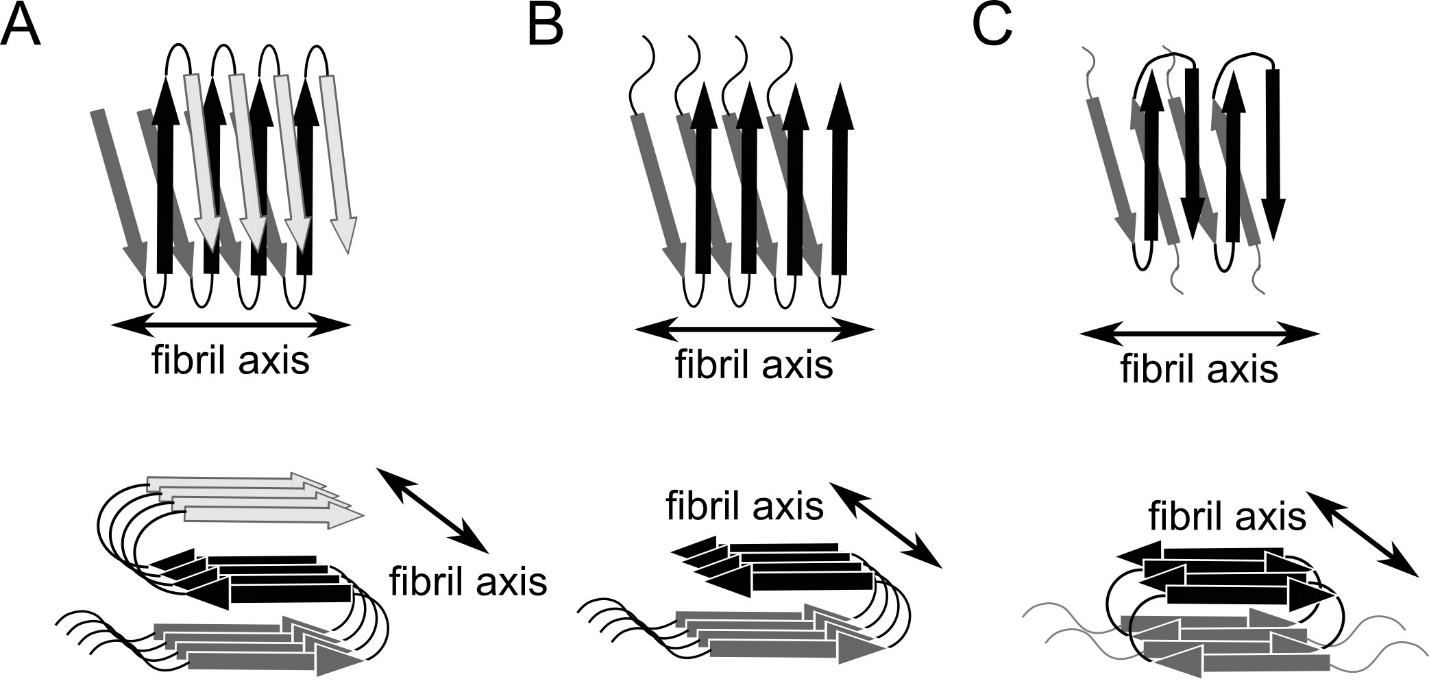

Figure S23. Diagrams of Aβ molecular conformation in different amyloid fibril structures. A) Aβ(1-42) fibril [6-8]. B) Aβ(1-40) fibril [9, 10]. C) Iowa mutant (D23N) of Aβ(1-40) fibril [11].
